# Epigenetic Remodeling Increases Plasticity of Bovine Fibroblasts Without Improving Embryo and Stem Cell Development

**DOI:** 10.64898/2026.10.28.762563

**Authors:** Dewison R Ambrizi, Diego Moreno, Juliano R Sangalli, Ricardo P Nociti, Jessica B Cruz, Laís V F Pessôa, David Kendall, Fabiana F Bressan, Marcos R Chiaratti, Lawrence C Smith, Pablo J Ross, Flavio V Meirelles, Rafael V Sampaio

**Affiliations:** Department of Veterinary Medicine, School of Animal Science and Food Engineering, University of São Paulo, Pirassununga, SP, Brazil; Department of Animal Science, University of California Davis, USA; Centre de Recherche en Reproduction et Fértilité, Faculty of Veterinary Medicine, University of Montreal, Saint-Hyacinthe, QC, Canada; Sexing Technologies, 22575 State Hwy 6 S, Navasota, TX 77868, USA; Department of Genetics and Evolution, Federal University of São Carlos, São Carlos, SP, Brazil; Department of Biomedical Sciences, Ontario Veterinary College, University of Guelph, Guelph, Ontario, Canada; Reproductive Sciences Unit, Toronto Zoo, Toronto, ON, Canada

**Keywords:** Epigenetic reprogramming, Chromatin remodeling, Somatic cell nuclear transfer, Small-molecule epigenetic modulators, Histone modifications, Bovine embryo, Cellular plasticity, Embryonic stem cells

## Abstract

Epigenetic memory, reinforced by redundant heterochromatic networks, is a major barrier to somatic cell reprogramming. In cattle, single-agent chromatin-modifying treatments show inconsistent efficacy, likely due to compensatory crosstalk among repressive marks. Here, we evaluated a multi-target small-molecule cocktail, GUSTC, combining <u>G</u>SK343 (EZH2 inhibitor), <u>U</u>NC0638 (G9a/GLP inhibitor), <u>S</u>odium Butyrate (HDAC inhibitor), <u>T</u>richostatin A (HDAC inhibitor), and <u>C</u>haetocin (SUV39H1 inhibitor) to simultaneously target facultative and constitutive heterochromatin in bovine fibroblasts. At optimized, non-cytotoxic concentrations, GUSTC treatment reduced H3K9me2, H3K9me3, H3K27me3, and HP1-alpha levels while increasing H3K9 acetylation by immunofluorescence. Using GUSTC-treated fibroblasts as donor cells for somatic cell nuclear transfer did not significantly change cleavage (83.6% vs. 92.2%; p = 0.15), blastocyst rate (28.4% vs. 39.0%; p = 0.20), bovine embryonic stem cell line derivation efficiency (26.3% vs. 33.3% of seeded blastocysts; p = 0.57), or the core pluripotency marker profile (OCT4, NANOG, SOX2) of the resulting lines. In a separate, serum-free, pluripotency-permissive culture system, GUSTC-treated cells adopted a colony-like morphology and showed increased *POU5F1* and *NANOG* expression, along with increased *DNMT1*, *EHMT2*, *HDAC1*, *HDAC2*, and *SUV39H1* transcripts. GUSTC thus reduces repressive heterochromatin marks in donor fibroblasts and is associated with a potentially more reprogramming-permissive state, without detectable improvement in preimplantation developmental competence. It provides a starting point for chemical reprogramming strategies in cattle.

**Summary sentence:** A small-molecule cocktail lowers H3K9me2/3, H3K27me3, and HP1-alpha in bovine fibroblasts and increases *POU5F1* and *NANOG* expression in serum-free culture, without improving cloned blastocyst development or embryonic stem cell establishment.

**Graphical abstract:** Proposed model: GUSTC treatment lowers repressive histone marks in bovine fibroblasts and, in serum-free culture, is associated with colony-like morphology and *POU5F1* and *NANOG* expression. In the model, GUSTC inhibits EHMT2, SUV39H1/2, EZH2, and HDACs, relieving repression and allowing H3K9 acetylation (AC); Me2/Me3, di-/trimethylation. In contrast, SCNT with GUSTC-treated donor fibroblasts did not significantly change cleavage, blastocyst development, or bESC derivation compared with control.

Alt text: Schematic titled "Proposed model". Left: timeline in which bovine fibroblasts in Alpha-MEM with 10% FBS are moved to KnockOut-DMEM/F12 with 20% KSR and GUSTC, form colony-like structures, and show increased plasticity. Center: in the top panel, EHMT2, SUV39H1/2, and the EZH2-containing PRC2 complex with HP1 add methyl groups to H3K9 and H3K27 while HDAC removes acetylation, so pluripotent genes are silenced; in the bottom panel, GUSTC blocks these enzymes, H3K9 is acetylated, and *POU5F1* and *NANOG* are on. Right: timeline in which fibroblasts in Alpha-MEM with 10% FBS plus GUSTC are used for nuclear transfer embryos and bovine embryonic stem cell colonies, with no significant change compared with control.

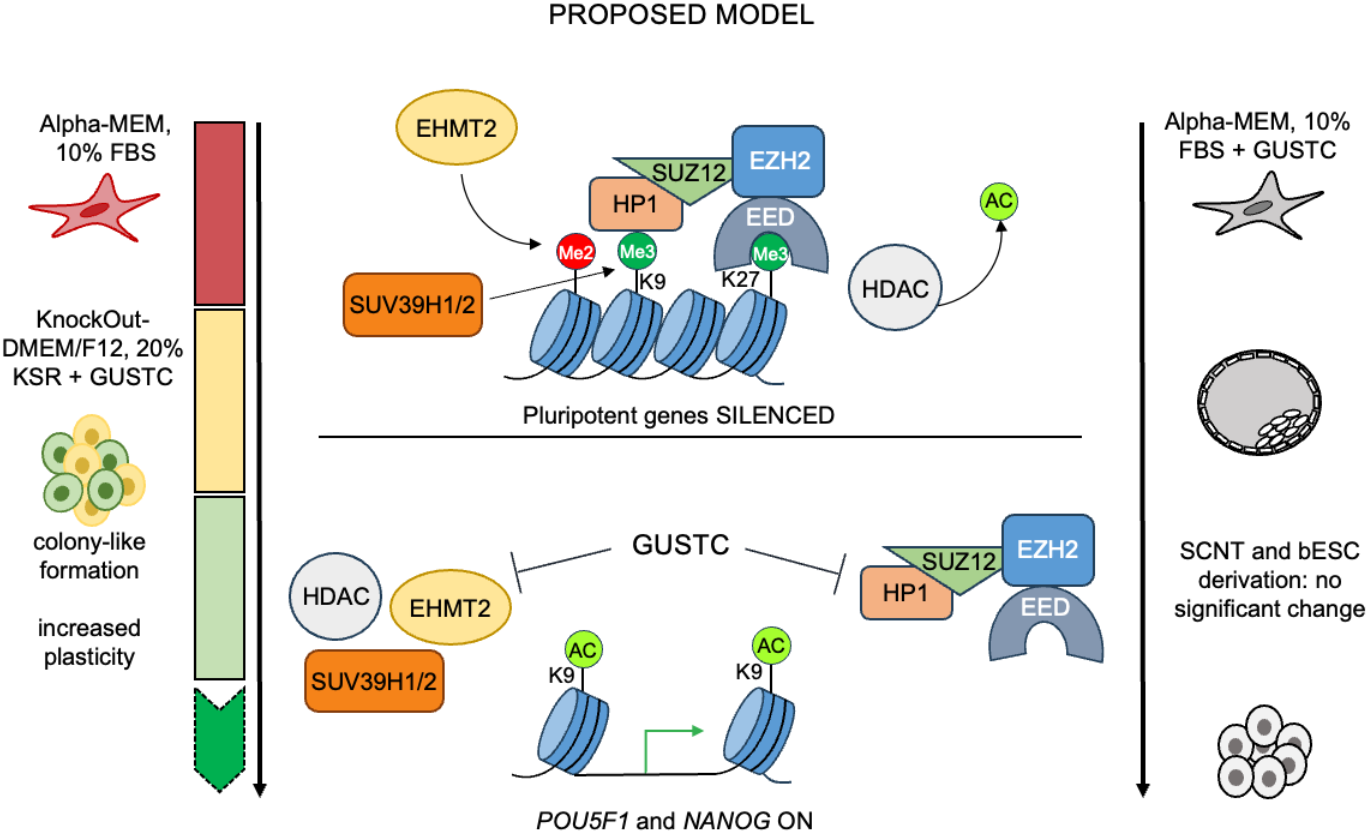

## Introduction

In higher eukaryotes, DNA and associated proteins assemble into chromatin, existing as either transcriptionally permissive euchromatin or densely repressive heterochromatin (1–5). Heterochromatin is further divided into facultative regions, marked by H3K27me3 and H3K9me2, and constitutive regions, stably repressed via H3K9me3 and HP1-alpha (6–8). This organization is established during development and mitotically propagated as a stable, functionally defining epigenetic memory (9), maintained through histone modifications and their interactions with chromatin regulators, including the Polycomb complexes PRC1 and PRC2, which together govern transcriptional output, nuclear organization, and promoter accessibility (10–21). In bovine embryos specifically, this memory is safeguarded by a small set of histone methyltransferases, EHMT1/2, SUV39H1/H2, SETDB1, and EZH2, responsible for depositing H3K9me2, H3K9me3, and H3K27me3, respectively (22), and is normally erased through the precisely timed, active expression of histone demethylases rather than passive replication-coupled dilution, a process that appears disrupted in cloned embryos, which display aberrant demethylation patterns (23,24).

Because of this tight regulation, histone and DNA hypermethylation constitute major barriers to nuclear reprogramming, limiting both the efficiency of somatic cell nuclear transfer (SCNT) and the generation of induced pluripotent stem cells (iPSCs) (25–28). A central obstacle is reactivating pluripotency genes that are stably silenced in somatic cells (29), which often resist even direct reprogramming cues such as the Yamanaka factors (30). Overcoming this resistance has therefore motivated strategies that combine targeted demethylation of repressive marks with reinforcement of permissive ones.

Small molecules that modulate chromatin marks are already used in cancer therapy and cellular reprogramming (31–35). Trichostatin A (TSA), for instance, has improved somatic cell reprogramming in mouse cloning (36), while sodium butyrate (NaBu) and 5-azacytidine have supported human iPSC generation (37). In cattle, however, these same treatments have largely failed to improve somatic cell reprogramming outcomes (38,39), an inconsistency that may reflect the extensive crosstalk among epigenetic marks, whereby altering one mark can trigger compensatory or unpredictable shifts in others (40). This crosstalk suggests that targeting repressive marks individually may be insufficient, and that a combinatorial strategy addressing multiple marks simultaneously could prove more effective.

GUSTC was designed to combine inhibitors acting on different steps in the establishment of repressive chromatin. GSK343 inhibits EZH2, the catalytic subunit of PRC2 that deposits H3K27me3 (32,40,41). UNC0638 is a selective chemical probe for the H3K9 methyltransferases G9a (EHMT2) and GLP (EHMT1), which deposit H3K9me2 (42); in donor cells, it has been reported to alter H3K9me2 and gene expression in cloned goat embryos (43). Chaetocin has been reported to inhibit SUV39H1, the methyltransferase that generates H3K9me3 (44), the mark bound by HP1-alpha (45). Trichostatin A and sodium butyrate inhibit histone deacetylases and thereby increase histone acetylation, a permissive mark (46,47). A combination of chaetocin and trichostatin A has also shown improved therapeutic effect in leukemia cells (35), supporting a combinatorial design. By acting on H3K27me3, H3K9me2/3, and histone acetylation together, GUSTC was expected to disrupt crosstalk among these marks rather than shift one.

Here, we tested this combinatorial hypothesis in bovine somatic cells using GUSTC, a cocktail of five small molecules: GSK343, UNC0638, sodium butyrate, trichostatin, and chaetocin, selected to simultaneously target the histone methyltransferases and deacetylases responsible for the repressive marks described above. We first characterized the effect of GUSTC on repressive chromatin marks in donor fibroblasts, and then asked whether this donor-cell epigenetic remodeling translated into improved developmental outcomes after nuclear transfer, by comparing cloning efficiency and bovine embryonic stem cell (bESC) line derivation between GUSTC- and control-treated donor cells. Finally, we examined whether GUSTC alters fibroblast plasticity in a serum-free, pluripotency-permissive medium, as a first step toward chemical reprogramming.

## Results

### Optimization of Chromatin-Modifying Molecule Concentrations

We followed a systematic approach to determine the optimal concentration of chromatin-modifying molecules for treating bovine fibroblasts. We selected the range of concentrations based on previous reports highlighting their effect on critical epigenetic marks for nuclear reprogramming (8,42). We studied how trichostatin (TSA), sodium butyrate (NaBu), chaetocin (CHTN), UNC0638 (UNC), and GSK343 (GSK) affect cell metabolism and morphology. Each molecule was tested alone by exposing fibroblasts to six different concentrations for 96 hours (Supplementary Figure S1). Complementary immunofluorescence-based dose–response for GSK343 confirmed a significant reduction in H3K27me3 at 100 nM relative to control (Supplementary Figure S2). The highest concentration that had minimal effect on cell viability, as assessed through an MTT assay, was selected for further experiments (indicated by red-shaded bars in Supplementary Figure S1). Then, we exposed cells to a combination of them at the recommended concentrations (GUSTC). Notably, these concentrations did not significantly affect cell viability, as demonstrated by only a 13% decrease in viability compared to the untreated group (Supplementary Figure S1). After treating cells with chromatin-modifying molecules, morphological changes were observed within 48–96 hours with some of the individual molecules (TSA and NaBu induced a filiform morphology; TSA increased and UNC0638 decreased nuclear size), but not with GUSTC (Supplementary Figure S3).

### Epigenetic Modulation by Chromatin-Modifying Molecules

To determine the impact of each chromatin-modifying molecule on epigenetic marks relevant to nuclear reprogramming, fibroblasts were treated individually with TSA, NaBu, CHTN, UNC0638, or GSK343, or with the combined cocktail (GUSTC), at the concentrations established above, for 96 hours, and H3K9me2, H3K9ac, H3K9me3, HP1-alpha, H3K27me3, 5mC, and 5hmC levels were measured by immunofluorescence. TSA, CHTN, and GSK343 increased H3K9me2, whereas NaBu, UNC0638, and GUSTC decreased it (Figures 1A, 1C, left). All treatments increased H3K9ac except NaBu, with GUSTC producing the largest increase by immunofluorescence (Figures 1A, 1C, right). H3K9me3 decreased with every treatment, most markedly with CHTN (Figures 1B, 1D, left), and HP1-alpha followed the same pattern (Figures 1B, 1D, right). H3K27me3 decreased with NaBu, UNC0638, and GSK343 but increased with TSA and CHTN (Figures 1E, 1G, left); GUSTC decreased H3K9me3, HP1-alpha, and H3K27me3 together (Figures 1B, 1D, 1E, 1G, left). For DNA methylation, TSA increased, and UNC0638 decreased 5mC (Figures 1F, 1H, left); 5hmC increased with NaBu and GSK343 and decreased with CHTN, while TSA and UNC0638 had no effect (Figures 1F, 1H, right). GUSTC had little effect on 5mC and increased 5hmC (Figures 1F, 1H). Western blotting confirmed that GUSTC decreased H3K9me2, H3K27me3, and HP1-alpha, with a non-significant trend toward lower H3K9me3 (p = 0.06), whereas the increase in H3K9ac observed by immunofluorescence was not confirmed (p = 0.15) and should be interpreted with caution (Figure 2; uncropped blots in Supplementary Figure S4). Principal component analysis of all measured marks separated GUSTC-treated from control fibroblasts into distinct clusters (PC1 and PC2 together explaining 58.1% of the variance; Supplementary Figure S5). Overall, a 96-hour exposure to TSA, NaBu, CHTN, UNC0638, or GSK343, alone or combined as GUSTC, altered multiple epigenetic marks without compromising cell viability.

**Figure 1.**
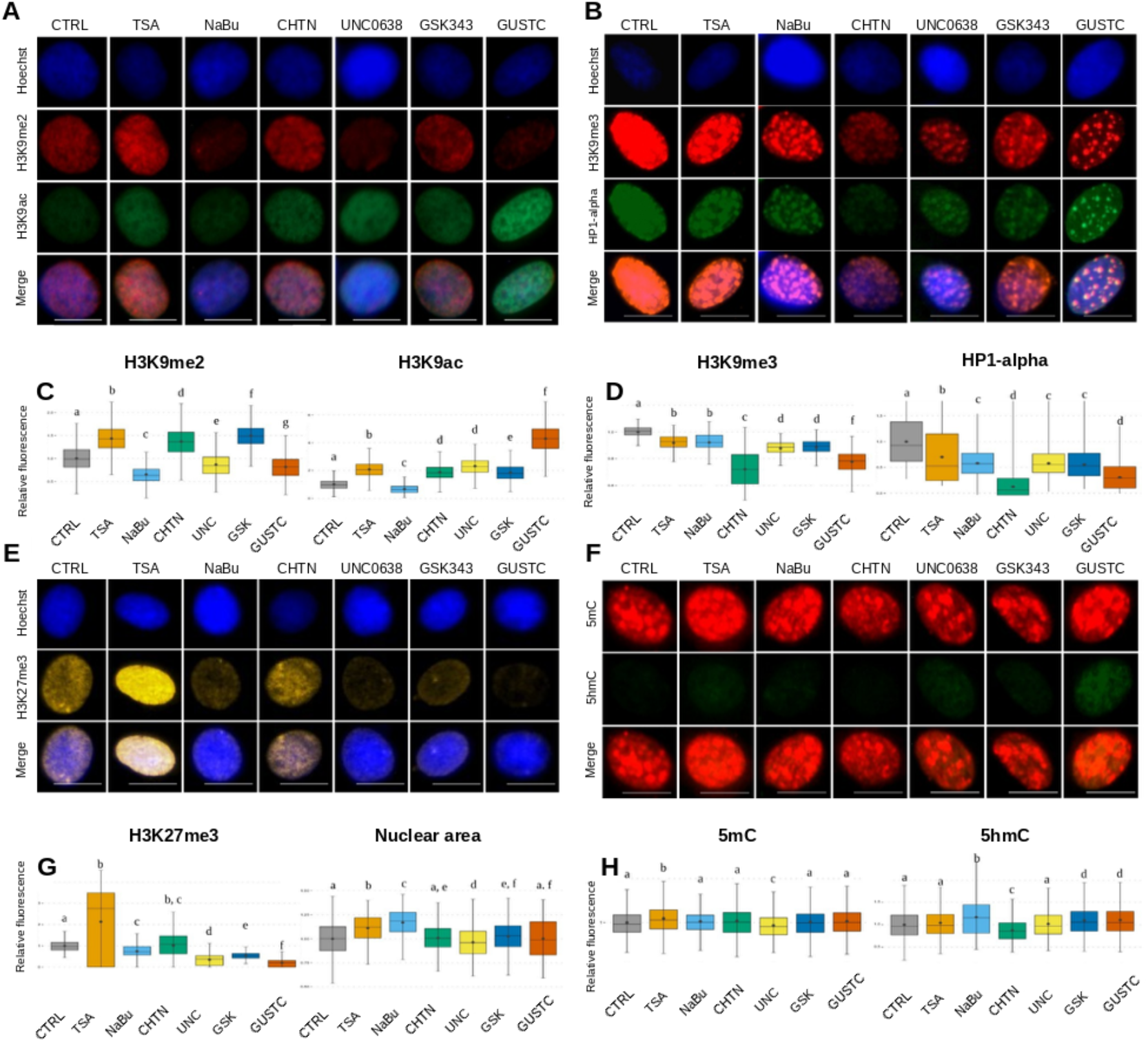
Immunostaining for repressive histone marks and DNA methylation in bovine fibroblasts treated with different chromatin-modifying molecules. (A) Representative nuclear immunostaining for H3K9me2 and H3K9ac, with treatments in columns and stains in rows; all images were acquired at the same magnification and exposure time (scale bar, 5 µm), and merged images are shown in the fourth row. (B) Representative nuclear immunostaining for H3K9me3 and HP1-alpha, with treatments in columns and stains in rows, under the same acquisition conditions as (A); merged images are shown in the fourth row. (C) Quantification of H3K9me2 (left) and H3K9ac (right) levels. (D) Quantification of H3K9me3 (left) and HP1-alpha (right) levels. (E) Representative nuclear immunostaining for H3K27me3, with treatments in columns and stains in rows, under the same acquisition conditions as (A); merged images are shown in the third row. (F) Representative nuclear immunostaining for 5mC and 5hmC, with treatments in columns and stains in rows, under the same acquisition conditions as (A); merged images are shown in the third row. (G) Quantification of H3K27me3 levels (left) and nuclear area, measured from Hoechst 33342 staining (right). (H) Quantification of 5mC (left) and 5hmC (right) levels. Groups not sharing a letter differ significantly (p < 0.05). On the x axes, UNC and GSK denote UNC0638 and GSK343, respectively. Data are from n = 3 wells per group, with approximately 800–1,000 nuclei analyzed per group; nuclei were the unit of observation (Kruskal–Wallis test followed by Conover test). Alt text: Eight-panel composite. Panels A, B, E, and F show fluorescence micrographs of single bovine fibroblast nuclei, with seven treatments in columns (CTRL, TSA, NaBu, CHTN, UNC0638, GSK343, and GUSTC) and stains in rows (Hoechst 33342 in panels A, B, and E): H3K9me2 and H3K9ac (A), H3K9me3 and HP1-alpha (B), H3K27me3 (E), and 5mC and 5hmC (F), each with a merged image. Panels C, D, G, and H show box plots of relative fluorescence for each mark (and nuclear area in G) across the seven treatments, with letters marking groups that differ significantly.

**Figure 2.**
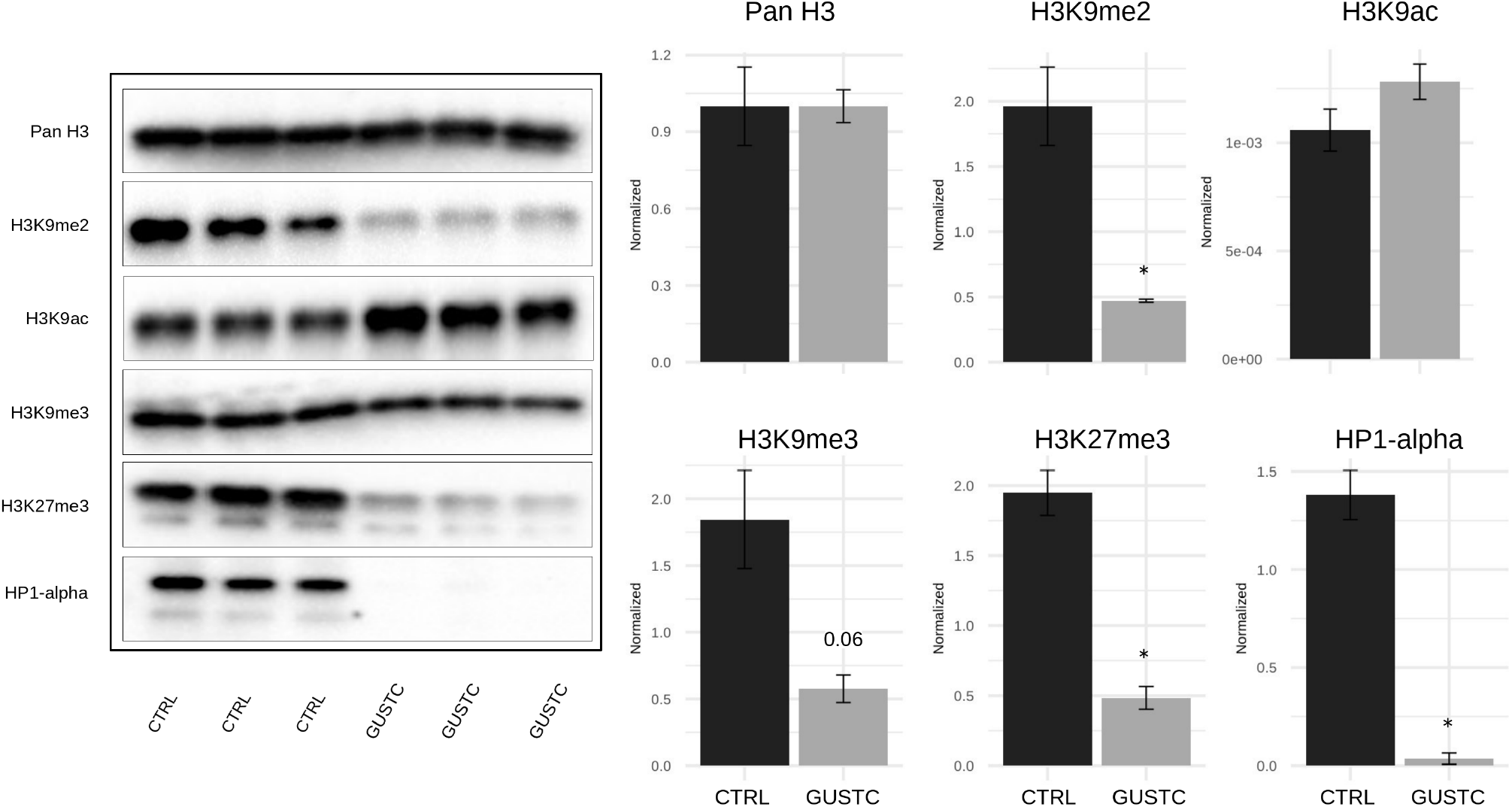
Western blot analysis of H3K9me2, H3K9me3, H3K27me3, HP1-alpha, and H3K9ac in control and GUSTC-treated bovine fibroblasts. Acid-extracted histones from day-4 cultures were analyzed by western blotting and probed in parallel using total histone H3 as a loading control; HP1-alpha was normalized to its own H3 loading control owing to differences in the extraction protocol (uncropped blots in Supplementary Figure S4). Statistical significance was set at p < 0.05. Histones were extracted from pooled cultures, and n = 3 lanes per group were quantified; the results therefore reflect technical rather than independent biological replication. Alt text: Western blot images of acid-extracted histones from three control (CTRL) and three GUSTC-treated fibroblast samples, probed for total H3, H3K9me2, H3K9ac, H3K9me3, H3K27me3, and HP1-alpha, with six bar graphs at right showing normalized band intensity for each target in CTRL versus GUSTC samples, with error bars. Asterisks mark significant differences, and one comparison (H3K9me3) is labeled p = 0.06.

### Cloning efficiency of GUSTC-treated donor cells

To determine whether the epigenetic remodeling observed in GUSTC-treated fibroblasts translated into improved developmental competence after nuclear transfer, we used GUSTC- or control-treated fibroblasts as donor cells for somatic cell nuclear transfer across five replicates (Table 1; replicate-level counts are provided in Supplementary Table S1). Donor cells were prepared in serum-containing medium (see Methods). Cleavage rates did not differ significantly between GUSTC (83.6%, 56/67) and control (92.2%, 71/77) embryos (Cochran–Mantel–Haenszel test stratified by replicate, p = 0.15). Blastocyst rates likewise did not differ significantly (GUSTC 28.4%, 19/67; control 39.0%, 30/77; p = 0.20). These results indicate that, despite the pronounced remodeling of repressive chromatin marks in donor fibroblasts, GUSTC treatment produced no detectable improvement in the developmental competence of the resulting cloned embryos. Because the 95% confidence intervals for these differences (Table 1) are wide, the data are compatible with a moderate reduction as well as with a small increase, and the study could detect only large differences (approximately 20 percentage points in blastocyst rate); the absence of a significant difference should therefore not be taken as evidence of equivalence.

**Table 1.**
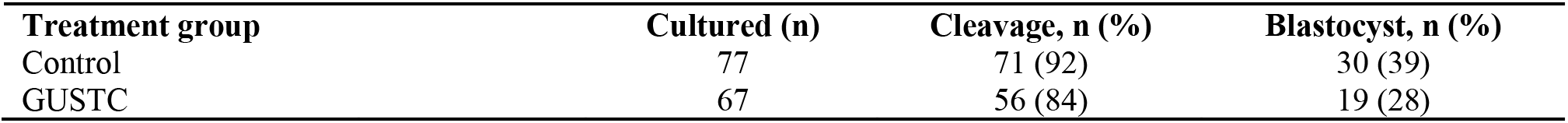
Cleavage and blastocyst rates of cloned embryos derived from GUSTC- or control-treated donor fibroblasts, pooled across five replicates. Replicate-level counts are provided in Supplementary Table S1. Differences (GUSTC minus control), Mantel–Haenszel estimates stratified by replicate: cleavage, −8.8 percentage points (95% CI, −19.1 to 1.4); blastocyst, −11.6 percentage *points (95% CI, −26.7 to 3.5)*.

| Treatment group | Cultured (n) | Cleavage, n (%) | Blastocyst, n (%) |
| --- | --- | --- | --- |
| Control | 77 | 71 (92) | 30 (39) |
| GUSTC | 67 | 56 (84) | 19 (28) |

### Bovine embryonic stem cell (bESC) line derivation from GUSTC- and control-derived clones

Blastocysts from both groups were seeded onto MEF feeder layers to assess whether donor-cell treatment affected bESC line establishment. bESC line derivation efficiency did not differ significantly between GUSTC- and control-derived clones, whether expressed relative to blastocysts seeded (26.3% vs. 33.3%) or to viable outgrowths (29.4% vs. 33.3%; Cochran–Mantel–Haenszel test, p = 0.57–0.82, depending on the denominator used). With 30 control and 19 GUSTC blastocysts seeded, only very large differences in derivation efficiency could have been detected. A session-by-session and passage-by-passage breakdown of derivation efficiency is provided in Supplementary Table S2.

### Statistical analysis of SCNT and bESC derivation outcomes

Cleavage and blastocyst rates, and bESC line derivation efficiency, were compared between GUSTC- and control-treated donor-cell groups using the Cochran–Mantel–Haenszel test stratified by replicate (Table 1, Supplementary Table S1) or by cloning session (Supplementary Table S2), performed in R (48). Between-group differences are reported as Mantel–Haenszel common risk differences (GUSTC minus control, in percentage points) with 95% confidence intervals (Greenland–Robins variance). With the sample sizes available, these analyses could detect only large differences; non-significant results are therefore interpreted from the confidence intervals and are not taken as evidence of no difference. Statistical significance was set at p < 0.05.

### Pluripotency marker expression in cloned bESC lines

To confirm the pluripotent identity of the derived lines, we characterized GUSTC- and control-clone bESC lines, along with a reference IVF-derived line, by alkaline phosphatase (AP) staining and immunofluorescence for SOX2, OCT4, NANOG, and SALL4 (Figure 3). All three lines (IVF, CLONE-Control, and CLONE-GUSTC) were AP-positive and expressed SOX2, OCT4, and NANOG at comparable levels, confirming that cloned lines from both treatment groups acquired a bona fide pluripotent phenotype indistinguishable from the IVF-derived reference. By contrast, SALL4 signal intensity appeared higher in the GUSTC-derived colonies than in the control-derived colonies (Figure 3C). To support this observation, we performed a semi-automated quantification of SALL4 immunofluorescence intensity from raw microscope files in bESC colonies from two independent derivation sessions per group (4 imaging fields per group in total, comprising 28 segmented colonies: 15 control-derived, 13 GUSTC-derived; representative fields and quantification shown in Supplementary Figure S6; field- and session-level values in Supplementary Table S4). Because the two sessions used different camera exposure times, net SALL4 signal was normalized to exposure time before comparison. As only two independent derivation sessions were available per group, the primary comparison was performed at the session level (n = 2 sessions per group), the most defensible unit of biological replication in this dataset: mean net SALL4 signal was higher in GUSTC-derived than in control-derived sessions in both sessions (∼2.1-fold on average; 1.9- and 2.3-fold in the two sessions), but this difference did not reach statistical significance given the small number of independent sessions (Welch’s t-test on session-level means, p = 0.234). A secondary, field-level comparison (n = 4 fields per group, of which 3 of 4 per group derive from the same session and are therefore not fully independent) was nominally significant (∼2.2-fold, p = 0.009). Because segmentation was automated and non-blinded, and the session-level comparison is likely underpowered at n = 2 per group, this result should be regarded as a hypothesis-generating trend rather than a confirmed finding, and a larger, formally replicated, and blinded quantification is needed.

**Figure 3.**
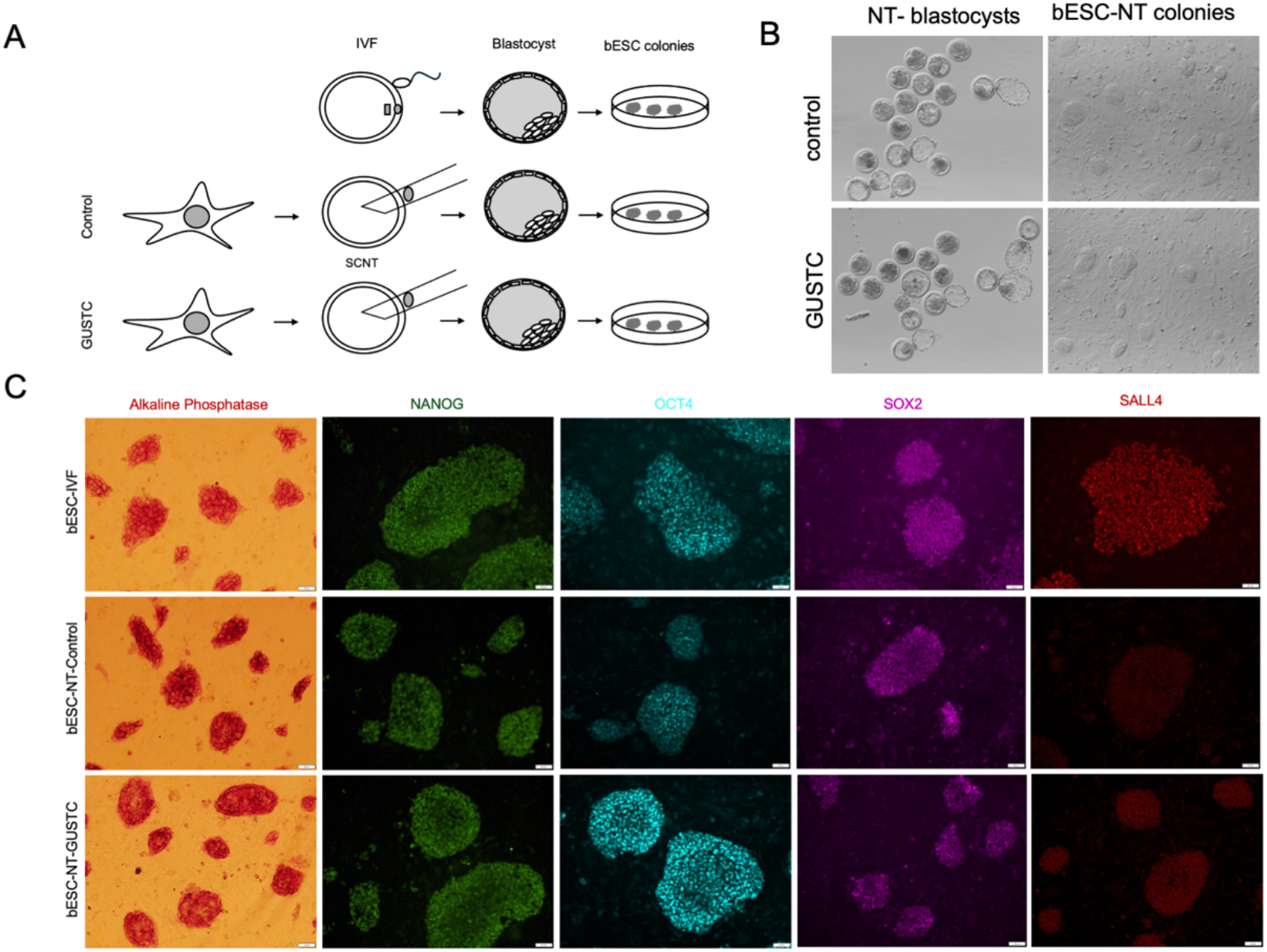
(A) Schematic overview of bovine embryonic stem cell (bESC) line derivation from in vitro fertilization (IVF)-produced embryos and from GUSTC- or control-treated somatic cell nuclear transfer (SCNT) donor cells. (B) Representative brightfield images of nuclear transfer (NT)-derived blastocysts and the resulting bESC-NT colonies from control and GUSTC-treated donor cells. (C) Pluripotency characterization of cloned bESC lines: Alkaline phosphatase (AP) staining and immunofluorescence for SOX2, OCT4, NANOG, and SALL4 in bESC lines derived from IVF embryos (reference) and from control- and GUSTC-treated cloned embryos. Cell lines and passages: for immunofluorescence, the IVF-derived line (passage 8), the control clone (passage 5), and the GUSTC clone (passage 6); for AP staining, the IVF-derived line (passage 10), the control clone (passage 7), and the GUSTC clone (passage 7). Scale bar in immunofluorescence images, 50 µm. Primary antibodies: SOX2, OCT4, and SALL4 (rabbit), NANOG (mouse); secondary antibodies: donkey anti-rabbit 568 and donkey anti-mouse 488. Exposure time and contrast were established from the IVF-embryo images and applied uniformly across all lines. The SALL4 signal of the GUSTC-derived colony appeared qualitatively closer in intensity to that of the IVF-derived reference colony than that of the control-derived colony; this three-way comparison is qualitative and was not part of the independent quantification shown in Supplementary Figure S6, which included only control- and GUSTC-derived material. Because SOX2, OCT4, and SALL4 were imaged in the same channel (donkey anti-rabbit 568), OCT4 and SOX2 are displayed pseudo-colored (cyan and magenta, respectively) for visual distinction; SALL4 is shown in its native acquisition color. Panel C shows one cell line per group (IVF-derived, control clone, and GUSTC clone). Alt text: Three-panel figure. (A) Schematic of bESC derivation from IVF embryos and from SCNT embryos made with control or GUSTC-treated donor fibroblasts, progressing from embryo to blastocyst to bESC colonies in a culture dish. (B) Brightfield images of nuclear transfer blastocysts and bESC-NT colonies for the control and GUSTC groups. (C) Grid of images for IVF-derived, NT-control, and NT-GUSTC bESC lines (rows) showing alkaline phosphatase staining and NANOG, OCT4, SOX2, and SALL4 immunofluorescence (columns); all three lines stain positive, and the SALL4 signal appears dimmer in the control clone than in the IVF and GUSTC lines.

### Shifting Cellular Plasticity through Epigenetic Modulation

Finally, we asked whether GUSTC treatment could alter fibroblast plasticity in a serum-free, pluripotency-permissive medium (KSR), as a first step toward chemical reprogramming. This assay used a different culture condition from the serum-containing protocol used to prepare SCNT donor cells, and these serum-free colony-forming cells were not used as SCNT donors, because higher *POU5F1* expression in donor cells has not improved the in vitro development of cloned bovine embryos, either in our previous studies (Sá et al. (49); Machado et al. (50)) or in the study by Goissis et al. (51). We measured expression of the pluripotency genes *FGF5*, *NANOG*, *POU5F1*, and *SOX2*, and of genes potentially responsive to chromatin-modifying treatment: *DNMT1*, *EHMT2*, *EZH2*, *HDAC1*, *HDAC2*, *KDM3A*, *KDM4A*, *KDM4C*, *KDM6B*, *SUV39H1*, *TET1*, and *TET2*. Fibroblasts were cultured with or without GUSTC for 96 h (day 4) and 192 h (day 8), and morphology was documented throughout (Figure 4A). Throughout the culture period, GUSTC produced no overt cytotoxic effects relative to control, although control fibroblasts proliferated visibly more slowly than GUSTC-treated cells under these conditions (Supplementary Figure S7). By 192 h, GUSTC-treated fibroblasts had adopted a colony-like morphology. *NANOG* expression increased at 192 h, and *POU5F1* expression increased at both 96 h and 192 h, relative to control (Figure 4B); *FGF5* and *SOX2* expression did not change significantly. Among the chromatin-modifying genes, *DNMT1*, *EHMT2*, *HDAC1*, *HDAC2*, and *SUV39H1* increased at 192 h, while *EZH2*, *KDM3A*, *KDM4A*, *KDM4C*, *KDM6B*, *TET1*, and *TET2* did not change significantly (Figure 4C). Together, these morphological and transcriptional changes, including increased expression of the pluripotency genes *NANOG* and *POU5F1*, are consistent with a shift toward a more plastic, potentially reprogramming-permissive state; however, pluripotency was not tested functionally.

**Figure 4.**
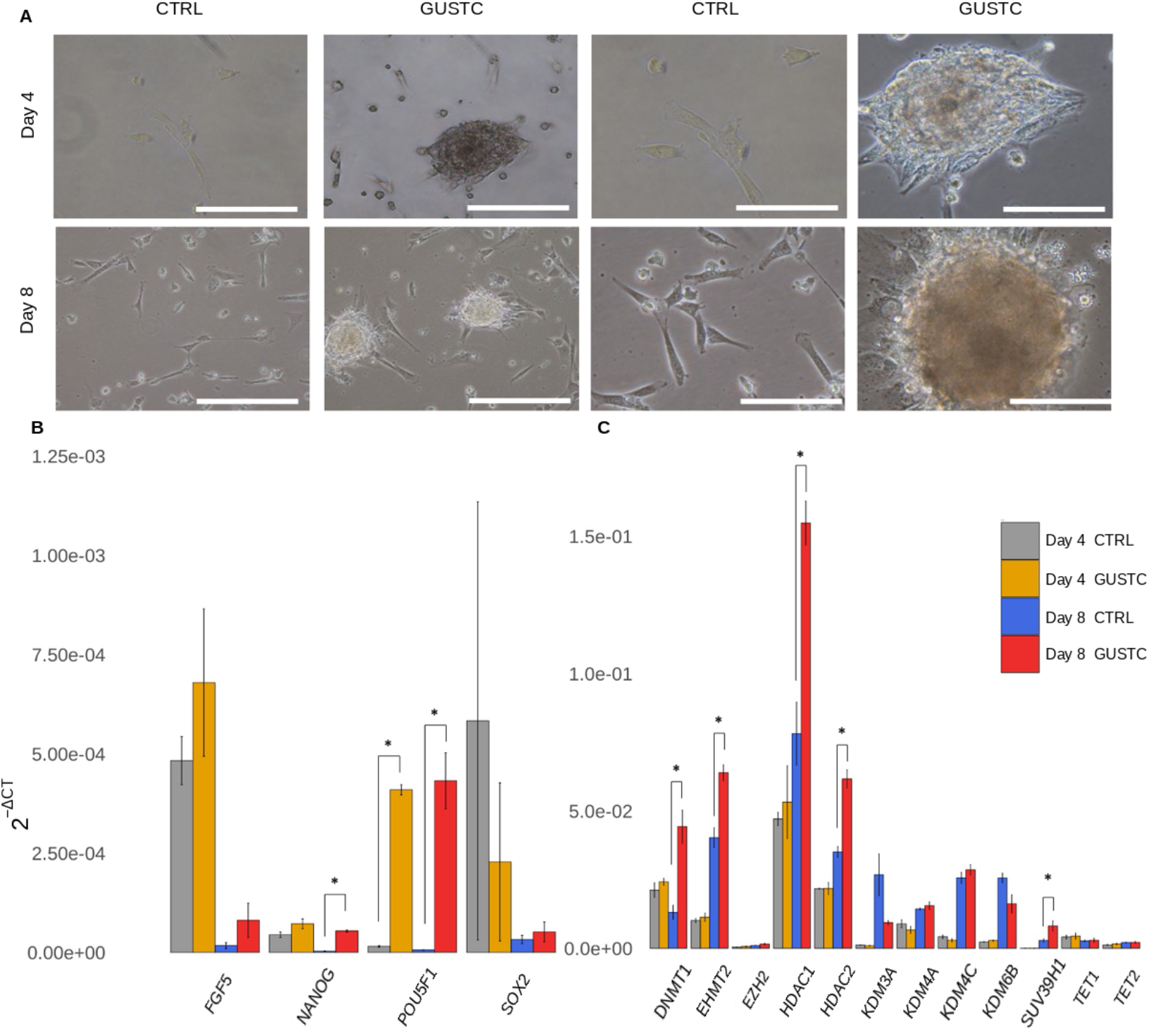
Morphology and gene expression of bovine fibroblasts cultured in serum-free, pluripotency-permissive medium. (A) Representative brightfield images of control and GUSTC-treated fibroblasts at days 4 and 8 (96 h and 192 h post-treatment, respectively); scale bars, 100 µm (left two columns) and 50 µm (right two columns). (B) Relative mRNA abundance of *FGF5*, *NANOG*, *POU5F1*, and *SOX2* (*NANOG* and *POU5F1* are normally repressed in somatic cells). (C) Relative mRNA abundance of epigenetic genes targeted directly or indirectly by the tested molecules. Expression is shown as 2^-ΔCT; error bars represent ± s.e.m. Asterisks indicate p < 0.05. n = 3 independent fibroblast lineages per group and time point. Alt text: (A) Brightfield micrographs of control and GUSTC-treated fibroblasts at day 4 and day 8; GUSTC-treated cultures contain dense cell aggregates that are not seen in controls. (B) Bar graph of relative mRNA abundance (2^-ΔCT) of *FGF5*, *NANOG*, *POU5F1*, and *SOX2* for control and GUSTC at day 4 and day 8, with asterisks marking significant differences for *NANOG* at day 8 and *POU5F1* at both times. (C) Bar graph of the same measure for *DNMT1*, *EHMT2*, *EZH2*, *HDAC1*, *HDAC2*, *KDM3A*, *KDM4A*, *KDM4C*, *KDM6B*, *SUV39H1*, *TET1*, and *TET2*, with asterisks for *DNMT1*, *EHMT2*, *HDAC1*, *HDAC2*, and *SUV39H1*.

## Discussion

GUSTC, a combination of five chromatin-modifying small molecules, remodeled the epigenetic landscape of bovine fibroblasts at low concentrations and without affecting cell viability, lowering the repressive marks H3K9me2, H3K9me3, H3K27me3, and HP1-alpha while increasing H3K9 acetylation by immunofluorescence (not confirmed by western blotting). Donor cells treated with GUSTC remained compatible with development to the blastocyst stage and with bESC derivation, with no significant difference from control donors, and the resulting lines expressed the core pluripotency markers OCT4, NANOG, and SOX2. Under serum-free, pluripotency-permissive conditions, GUSTC was further associated with a more permissive state, with a colony-like morphology and increased *POU5F1* and *NANOG* transcripts.

Low-concentration, extended exposure to these molecules, individually or combined as GUSTC, had minimal effect on cell metabolism by MTT assay (Supplementary Figure S1; GUSTC decreased it by approximately 13%) and, unlike some of the individual molecules, the combined treatment did not change cell morphology at 48 h, 72 h, or 96 h (Supplementary Figure S3), supporting the feasibility of these regimens for longer reprogramming protocols. H3K9 acetylation is a well-established permissive mark for chromatin accessibility and gene reactivation (52,53). TSA, an HDAC inhibitor, increased H3K9ac as expected; NaBu, also an HDAC inhibitor, did not (it lowered H3K9ac relative to the control), which may reflect a suboptimal concentration. CHTN, UNC0638, and GSK343 also increased H3K9ac, and the effect was largest with the combined GUSTC treatment by immunofluorescence. However, western blotting did not detect a significant change in H3K9ac with GUSTC (p = 0.15). This discrepancy may reflect the pooled, small-sample design of the western blots or differences between per-nucleus and bulk measurements, and it remains unresolved. H3K9 dimethylation is among the earliest-recognized epigenetic barriers to nuclear reprogramming (23), and several strategies have targeted this mark. In our previous work, targeting H3K9me2 did not improve SCNT outcomes in cattle (24). Here, NaBu was the most effective treatment for lowering H3K9me2, followed by GUSTC and UNC0638, a catalytic inhibitor of EHMT2, the enzyme that deposits this mark.

H3K9me3 was first shown to preserve epigenetic memory by suppressing key gene regions during SCNT in mouse (25), and in cattle, lower H3K9me3 is similarly associated with improved cloning outcomes (54). H3K9me3 increases during normal differentiation (55), whereas stem cells show lower H3K9me3 and more diffuse euchromatin (56,57). All compounds tested here decreased H3K9me3, most markedly CHTN, which inhibits SUV39H1 (the enzyme that deposits this mark), and GUSTC. Because CHTN also inhibits other histone lysine methyltransferases in vitro, including G9a, through covalent modification (58), part of its effect on H3K9 methylation may not be specific to SUV39H1, and overlap with the target of UNC0638 cannot be excluded. HP1 maintains heterochromatin integrity (59); mammals express three variants, alpha, beta, and gamma (60,61). Herein we focused on HP1-alpha because of its localization within constitutive heterochromatin (62) and because it is a useful indicator of heterochromatin-modulating interventions. HP1-alpha levels paralleled the H3K9me3 response across treatments, with the largest reduction seen with CHTN, consistent with the close mechanistic link between H3K9me3 and HP1 (16,44,63); GUSTC cocktail produced a similar reduction. The HDAC inhibitors NaBu and TSA reduced HP1-alpha, consistent with Taddei et al. (8); UNC0638 and GSK343 also reduced HP1-alpha, likely via their effects on H3K9me2 and H3K27me3, which are mechanistically linked to HP1 formation (17,45), consistent with human studies showing that small molecules can alter HP1 distribution (59,64).

The Polycomb group deposits H3K27me3, a mark linked to stable, heritable gene silencing (40). It is deposited by EZH2 and removed by KDM6A and KDM6B (65), and acts as an epigenetic barrier to nuclear reprogramming. Decreasing H3K27me3 marks in donor cells before reprogramming improves both SCNT and iPSC efficiency in pigs (66). In cattle, *KDM6A* overexpression was also related to H3K27me3 decrease and increased nuclear reprogramming (27). In our data, TSA increased H3K27me3 (Figure 1G, left), with greater variance than the other groups, suggesting phenotypic heterogeneity; this may relate to the concurrent increase in 5mC with TSA (Figure 1H, left), consistent with the reported correlation between H3K27me3 and 5mC (38,67,68). This contrasts with buffalo embryos, where TSA decreased H3K27me3 (69); the basis of this difference (species or cell type) is unclear, but it may reflect the crosstalk among chromatin marks proposed in multimeric-complex models (10,17). NaBu, UNC0638, GSK343, and GUSTC reduced H3K27me3, most markedly GUSTC, whereas TSA and CHTN yielded higher levels than the control (Figure 1G, left).

In this study, we did not include a dedicated 5mC inhibitor such as 5-aza-2′-deoxycytidine, but 5mC is important for heterochromatin maintenance and interacts closely with the other marks studied here (10,70–72). Overall, 5mC was more resistant to change than the other marks: only TSA and UNC0638 had modest effects on global 5mC. HDAC inhibitors have been linked to increased DNA methylation (73), and TSA has been reported to decrease *DNMT1* expression in cancer cells, particularly at high concentrations (74). DNMT1 interacts directly with EHMT2 (71), suggesting that UNC0638-mediated EHMT2 inhibition could have an effect on 5mC. We previously showed that EHMT1/2 inhibition alters 5mC and 5hmC dynamics during early bovine embryo development (24). 5hmC, which is generated from 5mC by TET enzymes (75,76), would be expected to decrease with lower 5mC levels; however, 5hmC was unchanged despite 5mC changes with TSA and UNC0638. On the other hand, NaBu, GSK343, and GUSTC increased 5hmC, while CHTN decreased it. Because the Polycomb complex is linked to both DNA methylation and H3K27me3, the 5hmC changes with GSK343 (an EZH2 inhibitor) may reflect redistribution of 5hmC following reduced H3K27me3 (77).

Despite this pronounced remodeling of repressive chromatin marks in GUSTC-treated donor fibroblasts, we detected no improvement in outcomes after nuclear transfer. Blastocyst rates (GUSTC 28.4% vs. control 39.0%; Cochran–Mantel–Haenszel test stratified by replicate, p = 0.20) and bESC line derivation efficiency (GUSTC 26.3–29.4% vs. control 33.3%; p = 0.57–0.82) did not differ significantly between groups. These estimates are imprecise (95% confidence intervals for cleavage and blastocyst rates are given in Table 1), and only large differences could have been detected; our data therefore show no improvement under the conditions tested but cannot exclude a modest effect in either direction. One interpretation is that remodeling of the repressive marks measured here in the donor cell prior to nuclear transfer is not, by itself, sufficient to overcome the additional epigenetic and developmental barriers encountered during preimplantation development. However, we did not measure these marks in reconstructed embryos, so this remains a hypothesis, and complementary interventions at the embryo stage may be needed.

One exception to this overall similarity was SALL4: in an exploratory quantification spanning two independent derivation sessions per group, mean net SALL4 immunofluorescence signal was higher in GUSTC-derived than in control-derived material in both sessions (1.9- and 2.3-fold), although the difference was not statistically significant at the session level (Supplementary Figure S6). Given the non-blinded, semi-automated nature of the analysis and the small number of independent sessions, this observation is hypothesis-generating only and should not be taken as evidence that GUSTC alters the pluripotency network of the derived lines. SALL4 is a core member of the pluripotency transcriptional network that acts alongside OCT4 and NANOG and is required for the specification of inner-cell-mass-derived lineages in the early embryo (78). In mouse embryonic stem cells, Sall4 binds thousands of target genes, forms a complex with Oct4 and Nanog, and its depletion lowers the expression of Oct4, Sox2, c-Myc, and Klf4 (79). Sall4 also enhances somatic-to-pluripotent conversion in cell-fusion assays (80), reconstitutes pluripotency from mouse fibroblasts together with Oct4 (81), and recruits DNA methyltransferases to silence non-pluripotency target genes (82). Whether the difference is reproducible, and whether it has any functional consequence, remains to be determined.

The absence of a detectable benefit of donor-cell remodeling is consistent with a central principle of nuclear reprogramming biology: after fusion, it is the oocyte cytoplasm, rather than the donor nucleus, that acts as the master regulator of reprogramming. Oocytes and eggs possess an intrinsically active reprogramming machinery, including an unusually high content of RNA polymerase II and the replication-independent histone variant H3.3, which drives rapid transcriptional reactivation of transplanted somatic nuclei largely independently of the donor cell’s prior chromatin state (83). Bovine oocytes and early embryos likewise express a defined repertoire of chromatin-remodeling and transcriptional-regulatory factors across the maternal-to-embryonic transition (24,84). Because somatic epigenetic memory reflects marks that accumulate in the differentiated state (85), pre- conditioning the donor cell, as achieved here with GUSTC, may lower the somatic memory entering the reconstructed embryo, but erasure of somatic identity and reactivation of the embryonic genome still depend on the oocyte’s own reprogramming machinery.

GUSTC, however, did alter fibroblast plasticity under permissive conditions in the absence of FBS. We hypothesized that fetal bovine serum might suppress reactivation of pluripotency genes, for example through serum components such as bone morphogenetic proteins (86), and tested this using a serum-free medium (see Methods). Under these serum-free conditions, GUSTC produced distinct morphological and transcriptional changes. This parallels findings from Guan et al. (87), who reprogrammed human fibroblasts to pluripotency using small molecules including epigenetic regulators: treated cells dedifferentiated, acquired epithelial characteristics, and formed OCT4-positive colonies that subsequently converted to chemically induced pluripotent stem cells. The morphological transformation and colony-like structures we observed in GUSTC-treated bovine fibroblasts are consistent with this framework, although further work is needed to determine whether these structures are transient and whether they express epithelial markers. In serum-free culture, GUSTC treatment also increased expression of *DNMT1* (Figure 4C). The same panel of repressive histone marks assessed in donor fibroblasts (H3K9ac, H3K9me2, H3K9me3, HP1-alpha, and H3K27me3) was evaluated by immunofluorescence under these conditions (Supplementary Figures S8–S10). Whether the increased *DNMT1* expression alters global 5mC remains to be determined, because 5mC was assessed only qualitatively in these cultures (Supplementary Figure S11). GUSTC also increased mRNA of *EHMT2* and *SUV39H1*, two methyltransferases that act on the same lysine residue. Because UNC0638 and CHTN inhibit these enzymes at the protein level, their increased transcription is consistent with a compensatory response in which the cell upregulates the chromatin writers targeted by GUSTC. *HDAC1* and *HDAC2* mRNAs were also upregulated alongside these methyltransferases, consistent with their cooperative role in gene silencing; some of these enzymes interact physically, which can enhance catalytic activity, as shown for the Sirt1-SUV39H1 interaction (88).

Pluripotency genes are normally repressed in somatic cells (30); our finding that GUSTC treatment was associated with increased OCT4/*POU5F1* and *NANOG* transcript abundance therefore represents a departure from this default state at the transcript level; whether chromatin remodeling at these loci underlies this change was not tested. Both genes are core components of the bovine pluripotency network and regulate embryo development and stem cell maintenance (89). OCT4/*POU5F1* is required for blastocyst development: its absence blocks progression beyond the morula stage and disrupts lineage specification (90). Nevertheless, neither high endogenous (49) nor exogenous (50,51) OCT4/*POU5F1* expression in donor cells improved in vitro development of cloned bovine embryos. This is of note because Goissis et al. reported that OCT4 expression in bovine donor fibroblasts reduced H3K9me3 and H3K27me3 (51), the same repressive marks that were lowered by GUSTC in the present study, and that OCT4 expression also increased the number of trophectoderm (CDX2-positive) cells in cloned blastocysts relative to YFP-expressing controls. OCT4/*POU5F1* is also required to maintain *NANOG* expression in both cattle and humans (91). *NANOG* itself is required for epiblast formation during the second lineage specification and for iPSC generation in cattle (92,93), making it a key determinant of pluripotency in this species. These findings suggest that GUSTC may create a more reprogramming-permissive state. Neither colony-like morphology nor increased *POU5F1* and *NANOG* transcript abundance demonstrates pluripotency acquisition; we did not assess protein expression, epigenetic resetting of pluripotency loci, or differentiation potential. In addition, proliferation was assessed only by visual inspection; the apparently faster expansion of GUSTC-treated cells relative to control (Supplementary Figure S7) was not quantified and could contribute to the differences in gene expression.

Because higher expression of *POU5F1* in donor cells did not improve the in vitro development of cloned bovine embryos in our previous studies (Sá et al. (49); Machado et al. (50)) or in that of Goissis et al. (51), we did not use the GUSTC-treated cells from the serum-free assay as nuclear donors; reactivation of *POU5F1* and *NANOG* transcription alone may not be sufficient. The plastic state described here is therefore a starting point for chemical reprogramming. Future work should define the factors and culture conditions that complete the acquisition of pluripotency (duration of culture, medium, gas atmosphere, and small-molecule composition) and, once cells with a stable, fully reprogrammed pluripotent phenotype are obtained, test them as nuclear donors.

This study has limitations. First, GUSTC combines five compounds acting on several chromatin marks, and individual agents changed some marks in opposite directions (for example, TSA and CHTN increased H3K27me3, whereas GUSTC decreased it). We compared single agents with the combination only for histone and DNA modifications in donor fibroblasts (Figure 1); SCNT, bESC derivation, and the serum-free assay used GUSTC alone, and we did not test synergy. The changes therefore cannot be attributed to any individual compound, and our data do not establish the molecular mechanism of GUSTC. Second, the SCNT and bESC outcomes rest on 67–77 cultured embryos per group across five replicates and 19–30 seeded blastocysts, so only large differences could be detected. Third, immunofluorescence was analyzed with nuclei as the unit of observation, western blot samples were pooled (technical replication), and gene expression was assessed at the transcript level only. Finally, colony-like morphology and increased *POU5F1* and *NANOG* transcripts are consistent with, but do not demonstrate, acquisition of pluripotency.

## Conclusion

Combined low-concentration treatment with the small-molecule cocktail (GUSTC) reduced the repressive marks H3K9me2, H3K9me3, H3K27me3, and HP1-alpha, and increased H3K9ac by immunofluorescence (not confirmed by western blotting), with the largest increase seen with GUSTC. However, this donor-cell epigenetic remodeling was not accompanied by a detectable improvement in blastocyst development or embryonic stem cell line derivation efficiency after nuclear transfer (the confidence intervals do not exclude moderate effects), suggesting that complementary interventions at the reconstructed-embryo level may be needed to translate donor-cell reprogramming into improved cloning outcomes. In a serum-free, pluripotency-permissive medium, by contrast, GUSTC-treated fibroblasts adopted a colony-like morphology and showed increased expression of the pluripotency genes *POU5F1* and *NANOG*, suggesting that heterochromatin in bovine fibroblasts, maintained through interdependent activity among KDACs, KDMs, DNMTs, and structural proteins such as HP1-alpha and histone H1, can be disrupted to yield a potentially more reprogramming-permissive state, although pluripotency was not tested functionally. These findings support GUSTC as a starting point for chemical reprogramming of bovine somatic cells; optimizing this approach toward complete reprogramming, and then testing the reprogrammed cells as nuclear donors, is the next step.

## Materials and Methods

### Animal care and use

All biological samples used in this study (donor fibroblasts and oocytes) were obtained post-mortem from cattle processed for commercial meat production at a local slaughterhouse, and the irradiated mouse embryonic fibroblasts used as feeder layers were purchased commercially. As no live animals were used or manipulated by the authors for the purposes of this research, approval by an institutional animal care and use committee was not required.

### Cell isolation

Bovine fibroblasts were obtained from a crossbred (*Bos taurus taurus* X *Bos taurus indicus)* calf. We collected skin post-mortem and minced the tissue with a scalpel blade. After this, the digestion was performed with 0.1% collagenase for three h at 38.5°C and 5% CO2 in a humid incubator. The digested material was centrifuged at 300 × g for 5 min, and the cell pellet was resuspended in culture medium alpha-MEM (Gibco BRL, Grand Island, NY, USA) supplemented with 10% Fetal Bovine Serum (FBS) (Gibco BRL) and 1% penicillin/streptomycin (Gibco BRL). The primary culture was grown in 60 mm Petri dishes until close to confluence. Cells were cultivated for two passages and then cryopreserved in cryotubes with a culture medium in addition to 10% dimethyl sulfoxide (DMSO) Sigma-Aldrich Chemical Company, (St. Louis, MO, USA) and stored at -80°C.

### MTT assay

The initial density was 1.5×10³ cells per well of 96-well plate (Costar Corning, Life Sciences). After 24h of initial culture, the medium was replaced with isolated molecules with six different concentrations for each of them. All molecules were bought from Sigma-Aldrich and diluted in DMSO with a stock concentration based on literature results. The concentrations were 1, 5, 10, 20, 40, 80 nM for TSA; 0.01, 0.05, 0.25, 1.5, 2 mM for NaBu; 1, 2.5, 5, 10, 20, 40 nM for CHTN; 0.01, 0.05, 0.25, 1.25, 6.25, 31.25 µM for UNC0638; 0.31, 0.62, 1.25, 2.5, 5 and 10 µM for GSK343. The control group (hereafter referred to as control throughout this manuscript) was supplemented with 1 µl/ml of DMSO alone. We tested time (48, 72, and 96 h) and concentration for each molecule in quintuplicate (plate wells). After the period cited, the culture media were removed, and then 90 µL of alpha-MEM serum-free containing 2.5mg/ml MTT (Sigma-Aldrich) remained for three h in the incubator. Then, 100 µL of DMSO was added to each well to solubilize the MTT formazan, and the absorbance was measured by the FLUOstar Optima microplate reader at 590 nm (BMG Labtech, Germany). The entire experiment was performed three times (n=3) for each molecule and time point. All statistical procedures were done in RStudio (48). Residue data were tested for distribution and then transformed using square-root transformation plus one. Using the Statistical Modeling package (94), we fit a linear mixed model with the plate as an uncontrollable factor. A post hoc test with Tukey adjustment evaluated differences among means. The values in the MTT analysis with all molecules in concomitant treatment at the same three-time intervals using the best concentrations were evaluated, and we performed the same analysis for the GUSTC group to estimate the decrease in cell metabolism. MTT graphs were constructed using ggplot2 (95) using the relativized value obtained by dividing the MTT value by the control MTT value. Error bars represent the standard error of the mean.

### Cell culture and treatment with chromatin-modifying molecules

Bovine fibroblasts were plated in a 24-well dish (Costar, Corning Life Sciences). Density was 15×10³ cells/well with alpha-MEM supplemented with 10% FBS and 1% penicillin/streptomycin. After 24 h of initial growth, media were prepared with isolated molecules or combined all of them in optimal concentrations based on the MTT assay: 10 nM for TSA, 0.25 mM for NaBu, 5 nM for CHTN, 0.25 µM for UNC0638, and 100 nM for GSK343. The medium was replaced daily, and cells were trypsinized on the second day of treatment.

### Cell immunofluorescence and analysis of epigenetic mark levels

For each group and modification, fibroblasts were cultivated in three different wells. After 96 h of treatment, all cells were fixed using ice-cold 100% methanol for 10 min at -20°C. We then washed them with PBS 3 times and stored them until the immunofluorescence day. For histone modifications and HP1, we proceeded as described previously (24,96) with few changes. We performed all immunofluorescence steps in a shaker, and except for the H3K9ac (Sigma H9286) antibody, we obtained the histone modifications and HP1 from Abcam. Briefly, cells were permeabilized using PBS + 1% Triton X-100 + 0.5% BSA for 12 min. For 5mC (ab10805) and 5hmC (ab214728) marks, we added an extra step: treat with deionized water with 4N HCl for 15min, then neutralize with Tris-Base (pH 8.8), as described previously (24). Then, in PBSW (PBS + 0.4% of Triton-X-100 + 0.1% Tween + 0.5% BSA) each antibody was diluted in concentrations close to those recommended by the manufacturer and then incubated for eight h at 4 °C. anti-H3K9ac was diluted in 1:1000; (ab1220) anti-H3K9me2 1:400; (ab8898) anti-H3K9me3 1:700; (ab6002) anti-H3K27me3 1:200; (ab77256) HP1-alpha 1:100. Wells were gently washed in PBSW 3 times for 10min. Afterward, secondary antibodies (Thermo Scientific, Waltham, MA, USA) against specific primary species were diluted in PBSW and incubated for one hour, protected from light. The cells were rewashed with PBSW for 10 minutes twice, followed by one more wash step with H33342 (2.5 µg/ml) for 10 minutes. The final wash step removed residual H33342. Immediately, wells were photographed using an epifluorescence microscope EVOS (Thermo-Fisher Scientific) using specific excitation light for each secondary antibody. Exposure time was standardized for each modification and was the same among groups (treatments). Three photos of each repetition, modification, and treatment were collected. We collected data with open-source software Fiji, using the DAPI filter as a guide and the tracking parameter for coordinates. The 5mC was used as a guide for DNA marks once the DNA denaturation made it impossible to counterstain with H33342. We tested residue distribution, then used the Kruskal-Wallis test followed by the Conover test because the residue distribution was non-normal. About 800 – 1000 nuclei were collected from each group.

### Acid extraction of histones from cells

Histones were extracted using the EpiQuick Total Histone Extraction Kit (Epigentek, Farmingdale, NY, USA, cat. # OP-0006) following the manufacturer’s instructions with a few modifications described for Sangalli et al. (97). Briefly, harvested cells were pelleted by centrifugation at 300 × g for 5 min at 4 °C. The cells were then incubated in 200 μL of pre-lysis buffer and centrifuged at 9,500 × g for 1 min at 4 °C. The supernatant was removed, and the pellet was diluted in 60 μL of lysis buffer and incubated for 30 min at 4 °C. Samples were centrifuged at 13,500 × g for 5 min at 4 °C. The supernatant, containing acid-soluble proteins, was transferred into a new vial and neutralized with 0.3 volumes of balance buffer supplemented with DTT. The proteins were quantified using Qubit 2.0 (Life Technologies) and then stored in aliquots at −20 °C until use.

### Coomassie staining

One microgram of acid-extracted histones was mixed with 4× Laemmli buffer (Bio-Rad Laboratories, Hercules, CA, USA, cat. #161-0747), denatured at 98 °C for 5 min, and loaded into an acrylamide gel. Proteins were fractionated by size on a 4%–15% SDS-PAGE gel run at a constant 100 V for 160 min. The gel was washed three times in ultrapure water and stained with SimplyBlue SafeStain (Thermo Fisher, cat. # LC6060) for one hour at RT. Next, the gel was washed for one hour in ultrapure water to remove the background. The gel image was captured on a ChemiDoc MP Imaging System (Bio-Rad), and histone enrichment was confirmed.

### Western Blotting for quantification of H3K9ac, H3K9me2, H3K9me3, HP1-alpha, and H3K27me3 levels in somatic cells

We prepared 0.5 μg of histones and ran them as described above to measure the relative levels. Next, the proteins were electroblotted onto PVDF membranes (Bio-Rad) at a constant 30 V for 70 min using a wet transfer system (Bio-Rad). The membranes were blocked with 5% BSA in Tris-buffered saline (TBS) + 0.1% of TWEEN (TBS-T) for one hour at room temperature. The membranes were incubated overnight at 4 °C under agitation with only one of the primary antibodies (anti-H3K9ac [Sigma, H9286], anti-H3K9me2 [Abcam, ab1220], anti-H3K9me3 [Abcam, ab8898], anti-HP1 [Abcam, ab77256] and anti-H3K27me3 [Abcam, ab6002]) diluted in concentration as recommended for manufacturer in TBS-T + 1% BSA solution. Subsequently, the membranes were washed three times with TBS-T for 5 min each and incubated for one hour with peroxidase-conjugated anti-rabbit secondary antibody (Sigma, A0545) diluted 1:5,000 in TBS-T + 1% BSA solution. The membranes were incubated for 1 min with Clarity Western ECL Substrate (Bio-Rad, 170-5060), and images were captured using the ChemiDoc MP Imaging System (Bio-Rad). Images were analyzed using the open-source software Fiji. The abundance of histone modification was calculated compared with loading control histone H3. Membranes were subsequently incubated overnight at 4 °C under agitation with an anti-histone H3 antibody (Sigma, H0164) diluted 1:10,000 in TBS-T + 1% BSA solution as a loading control, then washed and developed as described above; bands were quantified using Fiji, and the abundance of each histone modification was normalized to histone H3.

### Treated bovine fibroblasts cultured in serum-free, pluripotency-permissive medium with or without the molecules (plasticity assay)

This serum-free culture system, used exclusively to assess fibroblast plasticity and colony formation, is distinct from the serum-containing protocol used to prepare donor cells for somatic cell nuclear transfer (see “Somatic cell nuclear transfer and embryo culture”, below); fibroblasts cultured under these serum-free conditions were not used as SCNT donors. Bovine fibroblasts were plated in a 60 mm Petri dish and then placed in culture media to evaluate morphological alterations and evidence of increased epigenome plasticity. The medium was prepared as described by Bessi et al. (98) and consisted of KnockOut-DMEM/F12 (Gibco, 12660-012) supplemented with 20% KSR (Gibco, 10828-028), 1% GlutaMAX (Gibco, 35050-061), 1% MEM non-essential amino acids (Gibco, 11140-050), 1% penicillin-streptomycin (Gibco, 15140-122), 3.85 µM β-mercaptoethanol (Gibco, 21985-023), and 10 ng/mL bFGF (Peprotech, 100-18B). Then, the media were supplemented with the cocktail of epigenetic reprogramming molecules (GUSTC group) or not (Control group). Cells were cultivated for four and eight days in the presence or absence of the molecules across three different lineages, and morphology was additionally followed up to day 21 (Supplementary Figure S7).

### Gene expression analysis in fibroblasts

Bovine fibroblasts were cultured and treated with molecules, as described previously. Briefly, triplicate (n=3) 60mm plates were detached using Tryple Express (Gibco BRL), and the cell pellet was washed in PBS. Then, using RNeasy Plus Micro Kit (Qiagen, Hilden, Germany) and, following the protocol according to the manufacturer and adding 1 U of DNase I solution (Invitrogen, 18068015) for genomic DNA degradation, the total RNA was extracted and quantified using Nanodrop 2000 (Thermo Scientific). The RNA was converted to cDNA using a High-Capacity RNA-to-cDNA Kit (Life Tech, 4387406) using the manufacturer’s guidelines and stored at -20°C until use. We used the target genes listed in Supplementary Table S3 and the reference genes (*PPIA* and *RPL15*; Supplementary Table S3), as recommended by Livak (99). Primers used in RT-PCR were designed using Primer-BLAST (NCBI) based on sequences available in GenBank. The relative gene-specific mRNA quantification was performed using 20 µL reactions containing 0.2 mM of each primer plus 1 X Power SYBR Green PCR Master Mix (Life Technologies, 4367659) and 2 µL 8-fold diluted cDNA. All gene-specific cDNA-amplified samples were run in duplicate on the same qRT-PCR plate containing a negative control with 2 uL of ultrapure water instead of cDNA. The following cycling conditions were applied for amplification: initial denaturation at 95 °C for 10 min followed by 40 cycles of 95 °C for 15 seconds and 60 °C for 1 min. The SYBR Green fluorescence was read at the end of each extension step (60 °C). We confirmed amplification of a single product by analyzing melting curves and relative abundance using the comparative CT method (99). Relative abundance is expressed as 2^-ΔCT. ΔCT values of GUSTC-treated and control fibroblasts were compared for each gene and time point using a t-test (n = 3 independent fibroblast lineages per group); no correction for multiple comparisons was applied across genes, and these results should therefore be considered exploratory. Graphics were constructed using ggplot2 (95), and error bars represent the standard error of the mean.

### Somatic cell nuclear transfer and embryo culture

Somatic cell nuclear transfer was performed as previously described by Sampaio et al. (24), using donor fibroblasts treated with GUSTC or control for 4 days in alpha-MEM supplemented with 10% FBS and antibiotics prior to cloning. The only deviation from that protocol was the embryo activation procedure, which followed Felmer and Arias (100): reconstructed oocytes were activated with ionomycin (5 µM, 5 min), followed by anisomycin (1 µg/mL) and cytochalasin B (5 µg/mL) for 5 h. Cleavage and blastocyst development were recorded on the indicated days post-activation.

### Bovine embryonic stem cell (bESC) derivation and characterization

Blastocysts were seeded onto irradiated CF1 mouse embryonic fibroblast (MEF) feeder layers (A34180, Gibco, Thermo Fisher Scientific) in NBFR derivation medium, as previously described by Soto et al. (101), and outgrowths were monitored for bESC-like colony formation (Supplementary Figure S12) and serially passaged. Alkaline phosphatase activity was assessed using a commercial kit (Stemgent). For immunofluorescence characterization, colonies were stained for SOX2, OCT4, and SALL4 (rabbit primary antibodies) and NANOG (mouse primary antibody), followed by donkey anti-rabbit 568 and donkey anti-mouse 488 secondary antibodies; we standardized exposure time and contrast against the corresponding control line for each staining set. Because SOX2, OCT4, and SALL4 share the same secondary antibody (donkey anti-rabbit 568) and were therefore acquired in the same channel, OCT4 and SOX2 images were digitally pseudo-colored (cyan and magenta, respectively) for display in Figure 3C only, to allow visual discrimination between markers; SALL4 is shown in its native acquisition color, as it is the marker used for quantification. Pseudo-coloring consisted of a uniform linear remapping of raw pixel intensity to the new color channels, applied identically across all three cell lines, and did not alter any pixel values used in quantitative analysis.

SALL4 immunofluorescence intensity was further quantified in an independent set of bESC colonies, using raw 16-bit microscope TIFF files (Python 3, scikit-image, SciPy, NumPy) rather than processed display images. Colonies were drawn from two independent bESC derivation sessions per group, comprising 4 imaging fields per group in total (1 field per group from the first session, 3 fields per group from the second session; representative fields shown in Supplementary Figure S6A); because the two sessions used different camera exposure times for the SALL4 channel, net SALL4 signal was divided by the field’s exposure time before comparison. Colonies were segmented from the Hoechst channel by Gaussian smoothing (σ = 2 px, applied for segmentation only) followed by Otsu thresholding, morphological closing, and exclusion of objects smaller than 15,000 px²; mean raw SALL4 intensity was measured within each colony mask and background-subtracted using the local extracolony median signal from the same field (Supplementary Figure S6B). Because the primary session-level comparison (n = 2 independent sessions per group) is based on a small number of independent biological replicates, and segmentation was automated and non-blinded rather than manually curated, we report this quantification as a preliminary, exploratory, hypothesis-generating estimate rather than a validated formal measurement. Field- and session-level source data are provided in Supplementary Table S4; colony-level values and raw microscope files are available from the corresponding author upon reasonable request.

## Acknowledgment

The authors thank the staff and students at the LMMD and STgenetics, in particular Sergio Campos and Mark Yaeger. Dewison Ricardo Ambrizi was supported by São Paulo Research Foundation - FAPESP, grant number #2018/15089-0; Rafael Vilar Sampaio was supported by São Paulo Research Foundation - FAPESP, grant number #2015/08807-6, #2018/15089-0, and #2015/25111-5. This work was granted by São Paulo Research Foundation – FAPESP, grant numbers #2013/08135-2, #2014/50947-7, #2021/09886-8, #2022/15066-6, and #2023/04558-8; National Council of Technological and Scientific Development (CNPq) grant number 465539/2014-9; and CAPES. The funders had no role in the study design, data collection and analysis, publication decision, or manuscript preparation. During preparation of this manuscript, the authors used Claude (Anthropic) to assist with copyediting and text revision. The authors reviewed and edited all AI-assisted output and take full responsibility for the content of this publication.

## Author Contributions

DRA: Methodology, Investigation, Formal analysis, Writing – original draft. DM: Conceptualization, Methodology, Investigation, Formal analysis, Writing – original draft. JRS: Methodology, Investigation. RPN: Investigation. JBC: Resources, Methodology. LVP: Investigation. DK: Resources. MRC: Resources. FFB: Supervision, Resources. LCS: Writing – review & editing. PJR: Resources, Writing – review & editing. FVM: Conceptualization, Supervision, Funding acquisition. RVS: Conceptualization, Supervision, Funding acquisition, Writing – review & editing.

## Data Availability

Field- and session-level source data for the SALL4 quantification are provided in Supplementary Table S4; colony-level measurements and raw microscope files are available from the corresponding author upon reasonable request. Replicate-level counts for the nuclear transfer and bESC derivation experiments are provided in Supplementary Tables S1 and S2. All other datasets generated and analyzed during the current study are available from the corresponding author upon reasonable request.

