## Supplementary material for "Epigenetic Remodeling Increases Plasticity of Bovine Fibroblasts Without Improving Embryo and Stem Cell Development": Supplmental Material

*Supplementary Table S1. Cleavage and blastocyst rates of cloned embryos derived from GUSTC- or control-treated donor fibroblasts, by individual replicate. Pooled totals are reported in Table 1 of the main text.*

| Replicate | Cultured (n) | | Cleavage, n (%) | | Blastocyst, n (%) | |
|---|---|---|---|---|---|---|
| | Control | GUSTC | Control | GUSTC | Control | GUSTC |
| R1 | 16 | 14 | 16 (100) | 13 (93) | 7 (44) | 6 (43) |
| R2 | 18 | 18 | 18 (100) | 18 (100) | 11 (61) | 6 (33) |
| R3 | 9 | 9 | 7 (78) | 5 (56) | 2 (22) | 2 (22) |
| R4 | 13 | 14 | 11 (85) | 11 (79) | 4 (31) | 3 (21) |
| R5 | 21 | 12 | 19 (90) | 9 (75) | 6 (29) | 2 (17) |
| **Total** | **77** | **67** | **71 (92)** | **56 (84)** | **30 (39)** | **19 (28)** |

Cleavage and blastocyst percentages are calculated relative to the number of embryos cultured in each replicate and treatment group.

*Supplementary Table S2. Bovine embryonic stem cell (bESC) line derivation efficiency from GUSTC- and control-derived clones, by session and by donor-cell passage at the time of cloning.*

#### A. Session-by-session breakdown

| Session (passage) | Group | n | Viable outgrowths | bESCs | % bESCs/n | % bESCs / viable |
|---|---|---|---|---|---|---|
| 1 | Control | 7 | 7 | 2 | 28.6 | 28.6 |
| | GUSTC | 6 | 6 | 3 | 50.0 | 50.0 |
| 2 | Control | 11 | 11 | 5 | 45.5 | 45.5 |
| | GUSTC | 6 | 4 | 1 | 16.7 | 25.0 |
| 3 | Control | 2 | 2 | 2 | 100.0 | 100.0 |
| | GUSTC | 2 | 2 | 1 | 50.0 | 50.0 |
| 4 | Control | 4 | 4 | 1 | 25.0 | 25.0 |
| | GUSTC | 3 | 3 | 0 | 0.0 | 0.0 |
| 5 | Control | 6 | 6 | 0 | 0.0 | 0.0 |
| | GUSTC | 2 | 2 | 0 | 0.0 | 0.0 |
| **Total** | **Control** | **30** | **30** | **10** | **33.3** | **33.3** |
| | **GUSTC** | **19** | **17** | **5** | **26.3** | **29.4** |

Values are n unless otherwise noted. Total row sums across all five sessions; percentages are recalculated from the pooled totals, not averaged across sessions.

#### B. Passage-by-passage breakdown

| Group | P2–P3 (%) | P3–P4 (%) | P4–P5 (%) | P5–P6 (%) | Total (n) |
|---|---|---|---|---|---|
| Control | 20 | 50 | 20 | 10 | 10 |
| GUSTC | 20 | 60 | 20 | 0.0 | 5 |

Percentages indicate the proportion of established bESC lines derived from donor cells at each passage range at the time of cloning, relative to the total number of lines established per group.

*Supplementary Table S3. Primer sequences used for real-time qPCR (reference genes, DNA/histone methylation-pathway genes, and pluripotency genes).*

| Gene | Forward primer (5'–3') | Reverse primer (5'–3') | AT (°C) | Source |
|---|---|---|---|---|

| | | | | |
|---|---|---|---|---|
| PPIA (ref. gene) | GGTCCTGGCATCTTGTCCAT | TGCCATCCAACCACTCAGTCT | 60 | [1] |
| RPL15 (ref. gene) | CAAACGCCCAGTTCCTAAGG | TCGAGCAAACTTGAGCTGGTT | 60 | [1] |
| DNMT1 | GCAGTACCAGCCCATCCT | GCGGGCAGCCACCAA | 60 | [1] |
| EHMT2 (G9a) | GTTTGCCCTCCAGCTCAAC | GTCCACACCATTGACACAGG | 60 | [1] |
| TET1 | TGCCTACTTGCAACTGTCTTGATC | TCTATCCTTACTGCATTTCCTTTTTG | 60 | [1] |
| TET2 | GGATACACCTGTCAAGACTCAGTATGA | GGACCTGCTCCTAGATGGGTATAA | 60 | [1] |
| KDM3A | AGTGGCCTGCAATAATGTACAAA | GAAAGACGAAGATTGTTTACATCC | 60 | [2] |
| KDM4A | TGCTTCTGTAGCCTTGACCG | GGTCAGGCAGAAATCGCAGA | 60 | [2] |
| KDM4C | CCCAAGCAGTTCTCAAGGGT | GCTTCCTGGGTGATCTTGTCA | 60 | [2] |
| KDM6B | GGGAGACTATCAGCGCCTTC | AGCGGTACACAGGGATGTTG | 60 | [3] |
| POU5F1 (OCT4) | CTGGAGAAAGACGTGGTCCG | GAGACCCAGCAGCCTCAAAA | 60 | [4] |
| SOX2 | ATGGGCTCGGTGGTGAAGT | TGGTAGTGCTGGGACATGTGA | 60 | [4] |
| NANOG | TGTGGAGGAGAGCACAGAGA | CCAGGTCTTCACCTGCTTGT | 60 | [4] |
| HDAC1 | TTACGACGGGGATGTTGGA | GGCTTTGTGAGGGCGATAGA | 60 | [2] |
| EZH2 | TTCATGCAACACCCAACACTT | ACTCCTTCGCTCCCTCCAAG | 60 | This study |
| SUV39H1 | AGATCTATGACCGCCAGGGT | GGAGGTTGGGGTCACAACTG | 60 | [2] |
| FGF5 | CTCCCACGAAGCCAATATGT | GGCTTTTTCTTTTCGGGAAC | 60 | [5] |
| HDAC2 | AGTGTGGTGCAGACTCCCTA | TTGTGTATCCACCTCCCCCA | 60 | This study |

AT, annealing temperature. References: [1] Sampaio RV, Sangalli JR, De Bem THC, Ambrizi DR, Del Collado M, Bridi A, et al. Catalytic inhibition of H3K9me2 writers disturbs epigenetic marks during bovine nuclear reprogramming. Sci Rep 2020;10:11493. https://doi.org/10.1038/s41598-020-67733-9. [2] Sampaio RV. Modificações epigenéticas da cromatina e sua relação com a reprogramação nuclear de bovinos. 2016. https://doi.org/10.11606/T.10.2016.TDE-27082015-114515. [3] Glanzner WG, Rissi VB, Macedo MP de, Mujica LKS, Gutierrez K, Bridi A, et al. Histone 3 lysine 4, 9, and 27 demethylases expression profile in fertilized and cloned bovine and porcine embryos. Biology of Reproduction 2018;98:742–51. [4] Recchia K, Wathikthinnakon M, Bressan FF, Freude K. Generation of bovine iPSCs from fetal fibroblasts for in vitro myogenesis and cultured meat. Front Nutr 2025;12:1562981. https://doi.org/10.3389/fnut.2025.1562981 (source of primer sequences for POU5F1, SOX2, and NANOG). [5] Kawaguchi T, Tsukiyama T, Kimura K, Matsuyama S, Minami N, Yamada M, et al. Generation of naïve bovine induced pluripotent stem cells using piggyBac transposition of doxycycline-inducible transcription factors. PLoS One 2015;10:e0135403. https://doi.org/10.1371/journal.pone.0135403.

*Supplementary Table S4. Source data for the quantification of SALL4 immunofluorescence intensity in bovine embryonic stem cell (bESC) colonies derived from control- and GUSTC-treated donor cells (Supplementary Figure S6).*

#### A. Field level

| Group | Session | Field | Colonies (n) | Net SALL4 (a.u./ms), field mean |
|---|---|---|---|---|
| Control | 1 | 1 | 2 | 1.33 |
| Control | 2 | 2 | 5 | 1.82 |
| Control | 2 | 3 | 4 | 1.43 |
| Control | 2 | 4 | 4 | 1.77 |
| GUSTC | 1 | 1 | 6 | 2.47 |
| GUSTC | 2 | 2 | 2 | 3.79 |
| GUSTC | 2 | 3 | 1 | 4.10 |
| GUSTC | 2 | 4 | 4 | 3.62 |

#### B. Session level

| Group | Session | Fields (n) | Net SALL4 (a.u./ms), session mean |
|---|---|---|---|
| Control | 1 | 1 | 1.33 |
| Control | 2 | 3 | 1.68 |
| GUSTC | 1 | 1 | 2.47 |
| GUSTC | 2 | 3 | 3.84 |

#### C. Group comparison

| Comparison | n per group | Control, mean | GUSTC, mean | Fold change | Welch's t-test, P |
|---|---|---|---|---|---|
| Session level (primary) | 2 | 1.51 | 3.15 | 2.1 | 0.234 |
| Field level (secondary) | 4 | 1.59 | 3.50 | 2.2 | 0.009 |

a.u./ms, arbitrary units per millisecond of SALL4-channel camera exposure. For each colony, net SALL4 signal was calculated as the mean raw intensity within the colony mask minus the median signal outside the colonies in the same field, divided by the exposure time of that field (the two sessions used different exposure times). Field values are the mean of the colonies within a field (28 colonies in total: 15 control-derived and 13 GUSTC-derived); session values are the mean of the field values within an independent derivation session. Fold change is the ratio of the GUSTC to the control group mean. P values are from two-sided Welch's t-tests on session means (primary comparison, n = 2 per group) and on field means (secondary comparison, n = 4 per group; three of the four fields per group derive from the same session and are therefore not fully independent). The session-level comparison is underpowered and the analysis is exploratory (automated, non-blinded segmentation). Colony-level values, raw microscope files, and the analysis script are available from the corresponding author upon reasonable request.

### Supplementary Figure Legends

**Supplementary Figure S1.** MTT assay dose-response for each chromatin-modifying molecule in bovine fibroblasts over 96 hours. Trichostatin A (TSA) showed no difference in metabolism relative to control at 10 nM; sodium butyrate (NaBu) increased metabolism at low concentrations but not at 0.25 mM; Chaetocin (CHTN) showed no difference at 5 nM; UNC0638 tolerated a wide concentration range before affecting metabolism, and 0.25 µM (consistent with prior reports) was sufficient to decrease H3K9me2 levels; GSK343 at 100 nM did not alter metabolism. The combined treatment (GUSTC) decreased cell metabolism by approximately 13% relative to control. Concentrations selected for further experiments are indicated by red-shaded bars.

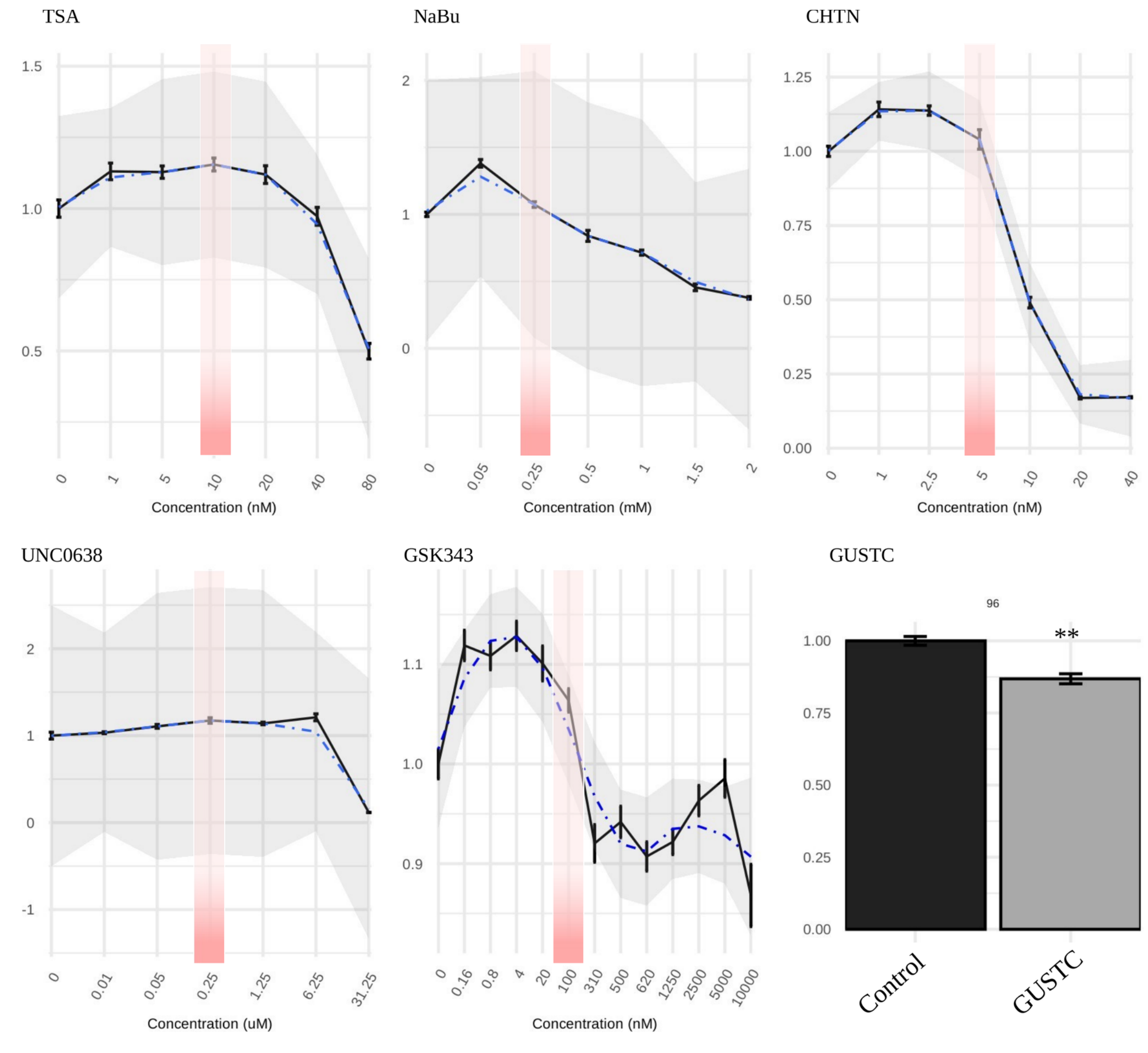

**Supplementary Figure S2.** Dose-response of GSK343 on H3K27me3 levels by immunofluorescence. At 100 nM, H3K27me3 fluorescence intensity was significantly reduced relative to control. Individual nuclei; scale bar, 5 µm.

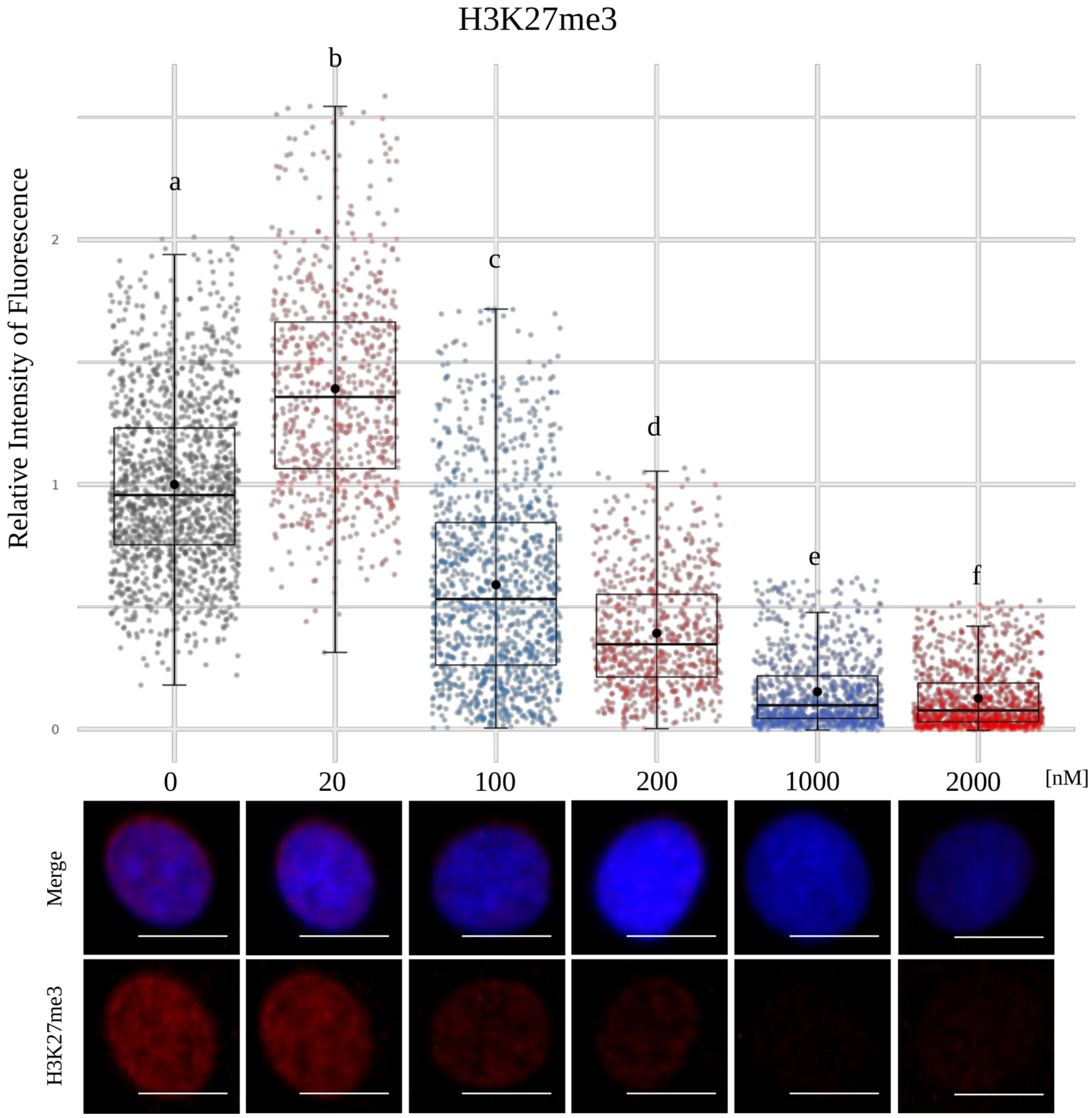

**Supplementary Figure S3.** Bovine fibroblast morphology at 48, 72, and 96 hours after treatment. No treatment reduced proliferation relative to control. TSA increased nuclear size and induced a filiform morphology by 96 hours; sodium butyrate increased proliferation and induced a filiform morphology by 72 hours. Chaetocin, UNC0638, and GSK343 showed no difference from control, although UNC0638 decreased nuclear size. GUSTC did not alter morphology or proliferation relative to control.

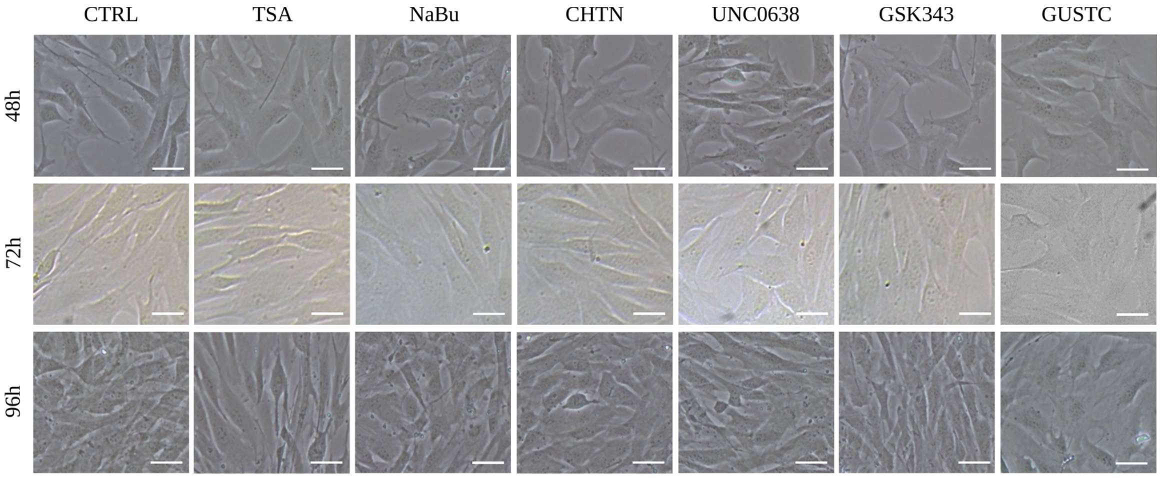

**Supplementary Figure S4.** Uncropped western blot images corresponding to Figure 2.

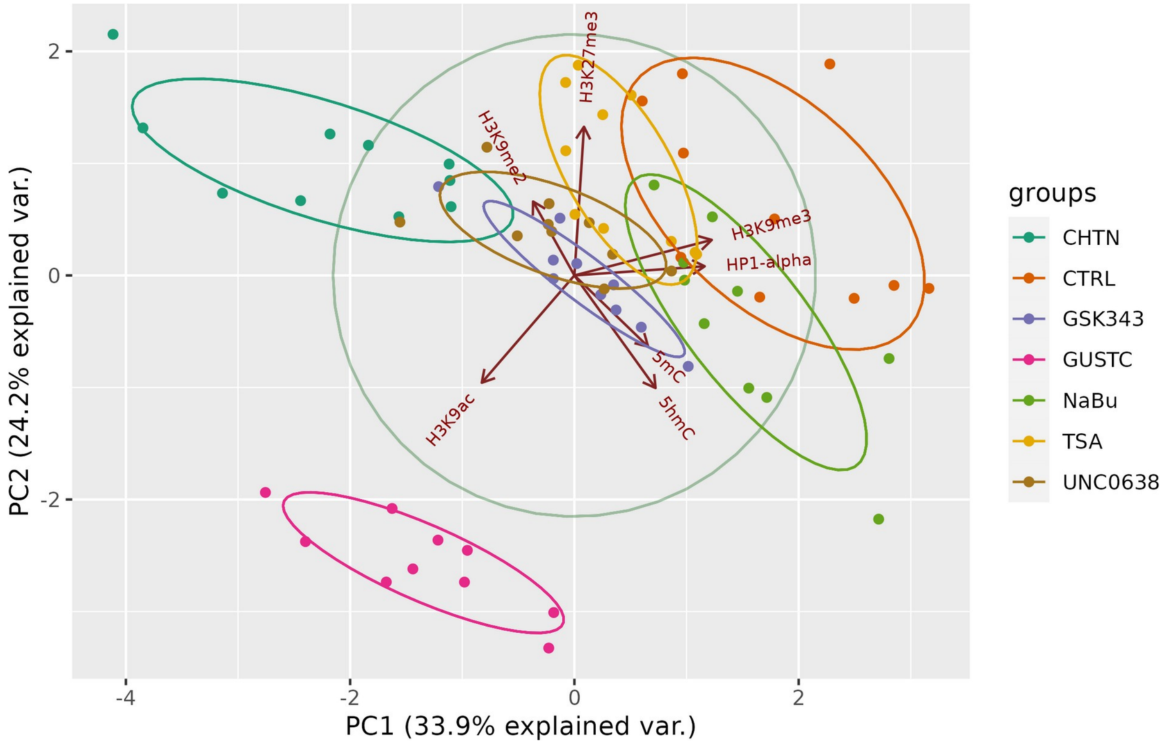

**Supplementary Figure S5.** Principal component analysis (PCA) of all measured epigenetic marks, grouped by treatment and replicate. PC1 and PC2 together explain 58.1% of the variance in the data; control (orange) and GUSTC (pink) groups formed distinct clusters, consistent with the treatment-associated differences described in the main text.

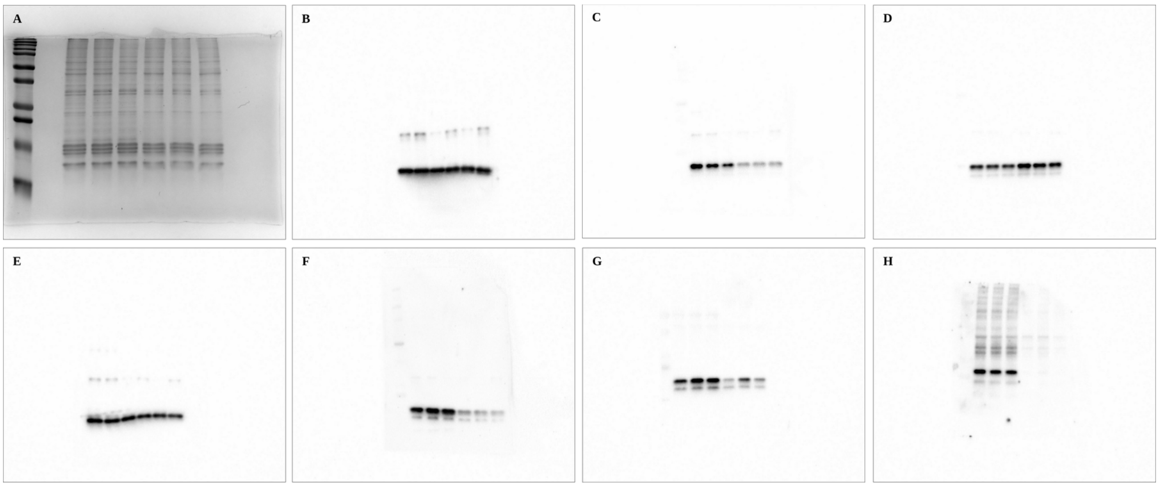

**Supplementary** Figure S6. Quantification of SALL4 signal intensity in an independent set of bovine embryonic stem cell (bESC) colonies from two independent derivation sessions per group. (A) Representative composite immunofluorescence images (SALL4, red; Hoechst, blue) from four imaging fields per group (1 field from the first session, 3 fields from the second session; exposure times differed between sessions and signal was exposure-normalized for quantification), for control- and GUSTC-derived colonies. Scale bars, 50 µm. (B) Net SALL4 intensity per colony, normalized to exposure time (background-subtracted; 28 colonies pooled from the fields shown in (A): 15 control-derived, 13 GUSTC-derived). Colonies were segmented from the Hoechst channel using Gaussian smoothing, Otsu thresholding, and morphological closing; boxes show the median and interquartile range, whiskers show the full range. At the session level ($n = 2$ independent sessions per group, the primary comparison), mean net SALL4 signal was higher in GUSTC-derived than control-derived material in both sessions (~2.1-fold on average) but did not reach statistical significance (Welch's t-test on session-level means, $p = 0.234$); a secondary, field-level comparison ($n = 4$ fields per group, partially non-independent) was nominally significant ($p = 0.009$; field- and session-level values are given in Supplementary Table S4). This quantification used raw microscope files but automated, non-blinded colony segmentation, and should be interpreted as a preliminary, hypothesis-generating estimate rather than a validated formal measurement.

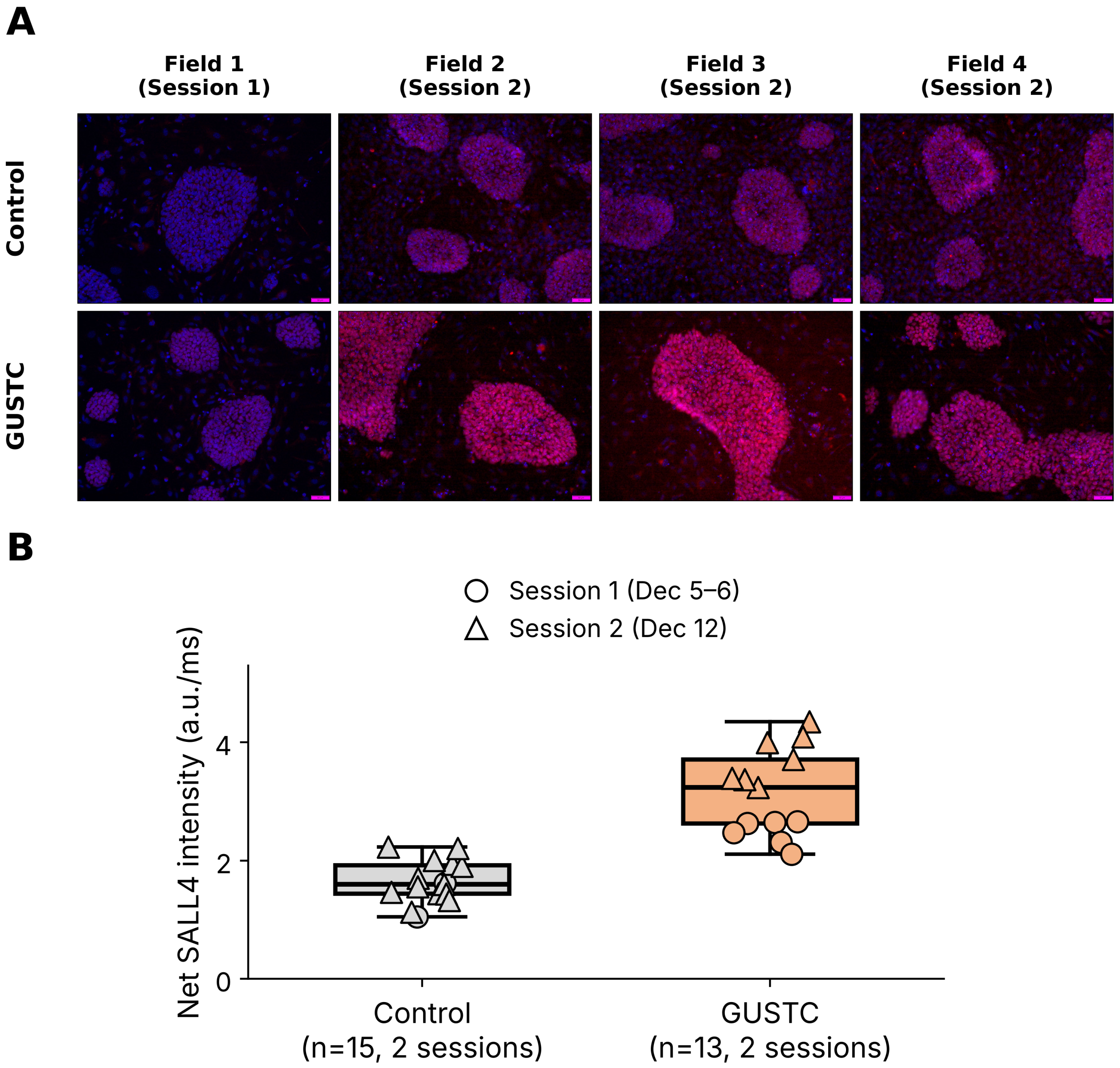

**Supplementary** Figure S7. Bovine fibroblast morphology in serum-free, pluripotency-permissive medium (KSR) from day 2 to day 21. Control and GUSTC groups are shown in columns (scale bars, 100 µm in the left two columns and 50 µm in the right two columns), with time shown in rows. Neither group showed overt cytotoxic effects; control fibroblasts proliferated more slowly than GUSTC-treated fibroblasts under these conditions.

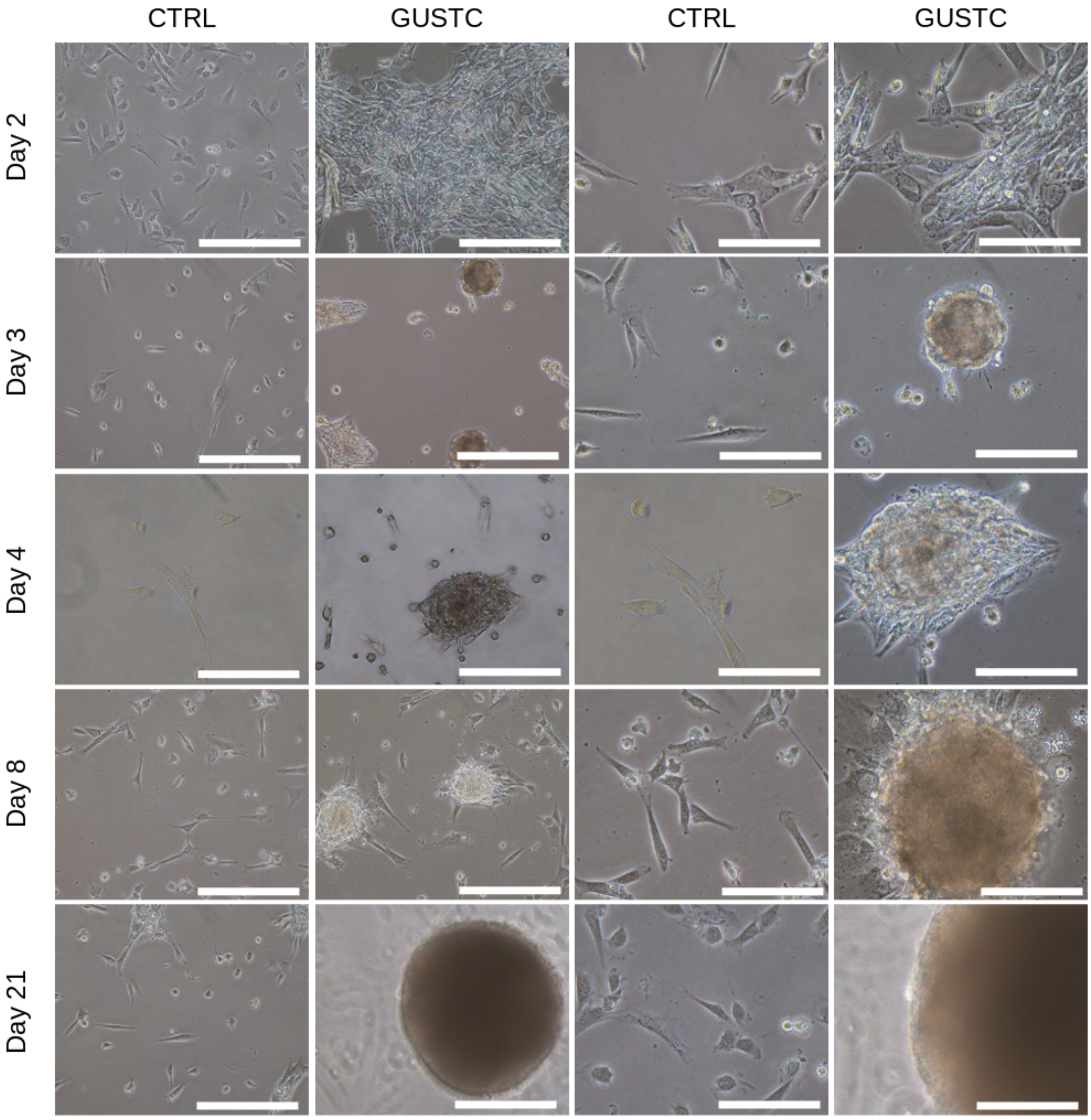

**Supplementary** Figure S8. Immunofluorescence for H3K9ac and H3K9me2 in fibroblasts cultured in serum-free, pluripotency-permissive medium, treated (GUSTC) or not (control) with chromatin-modifying molecules. Hoechst (blue), H3K9ac (red), H3K9me2 (green), and merge. All images were acquired at the same magnification and exposure time.

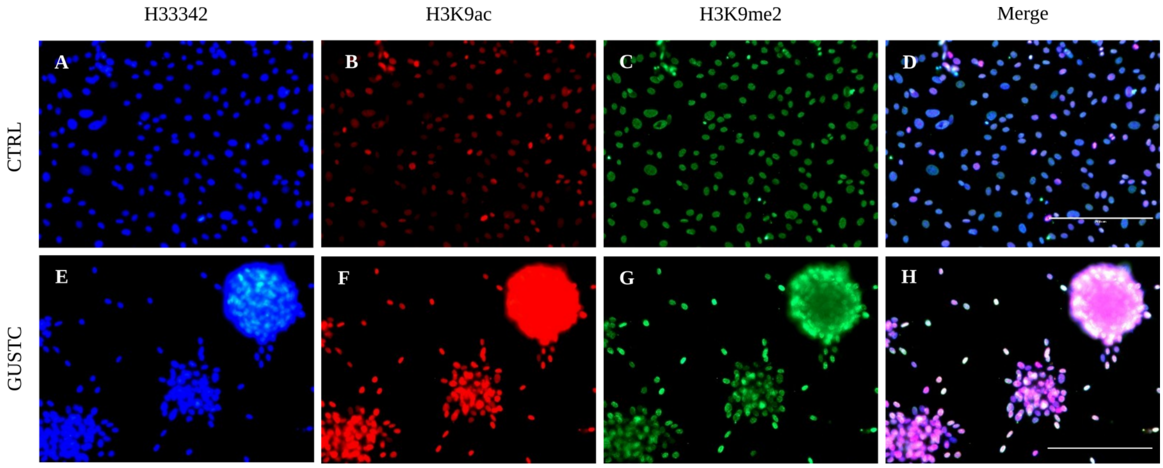

**Supplementary** Figure S9. Immunofluorescence for H3K9me3 and HP1-alpha in fibroblasts cultured in serum-free, pluripotency-permissive medium, treated (GUSTC) or not (control) with chromatin-modifying molecules. Hoechst (blue), H3K9me3 (red), HP1-alpha (green), and merge. All images acquired at the same magnification and exposure time.

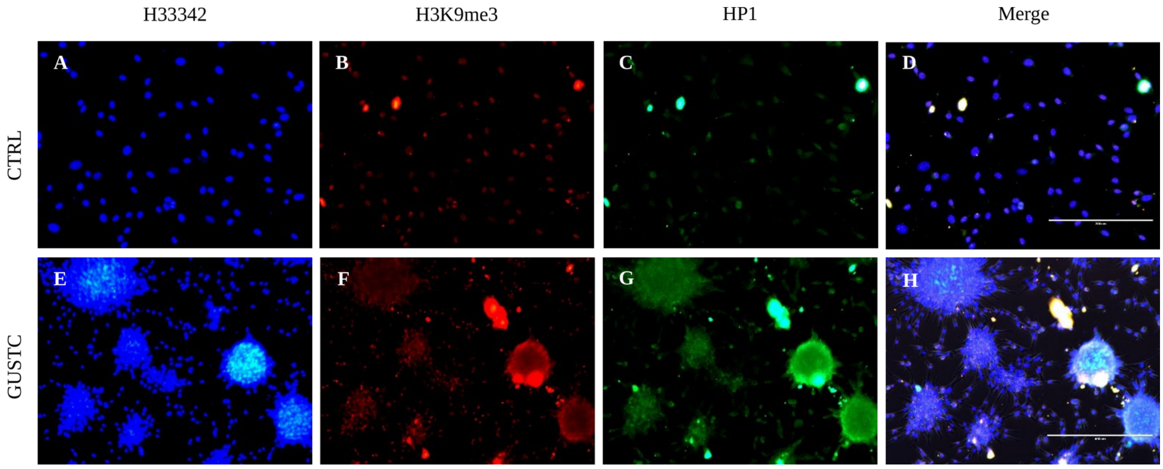

**Supplementary** Figure S10. Immunofluorescence for H3K27me3 in fibroblasts cultured in serum-free, pluripotency-permissive medium, treated (GUSTC) or not (control) with chromatin-modifying molecules. Hoechst (blue), H3K27me3 (green), and merge. All images acquired at the same magnification and exposure time.

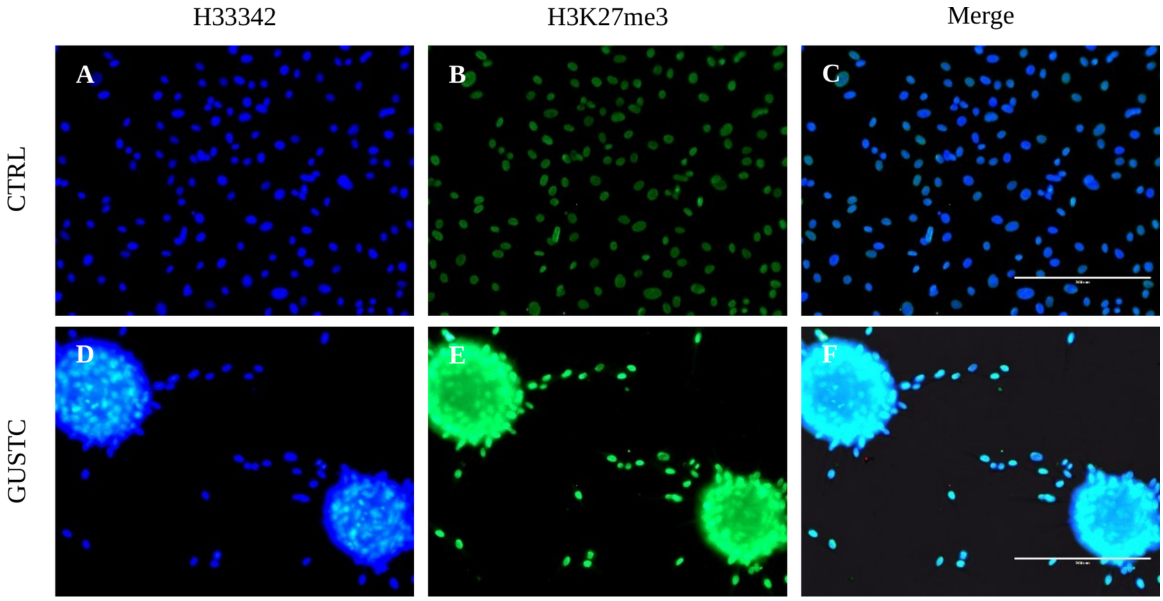

**Supplementary** Figure S11. Immunofluorescence for 5mC and 5hmC in fibroblasts cultured in serum-free, pluripotency-permissive medium, treated (GUSTC) or not (control) with chromatin-modifying molecules. 5mC (red), 5hmC (green), and merge. All images were acquired at the same magnification and exposure time.

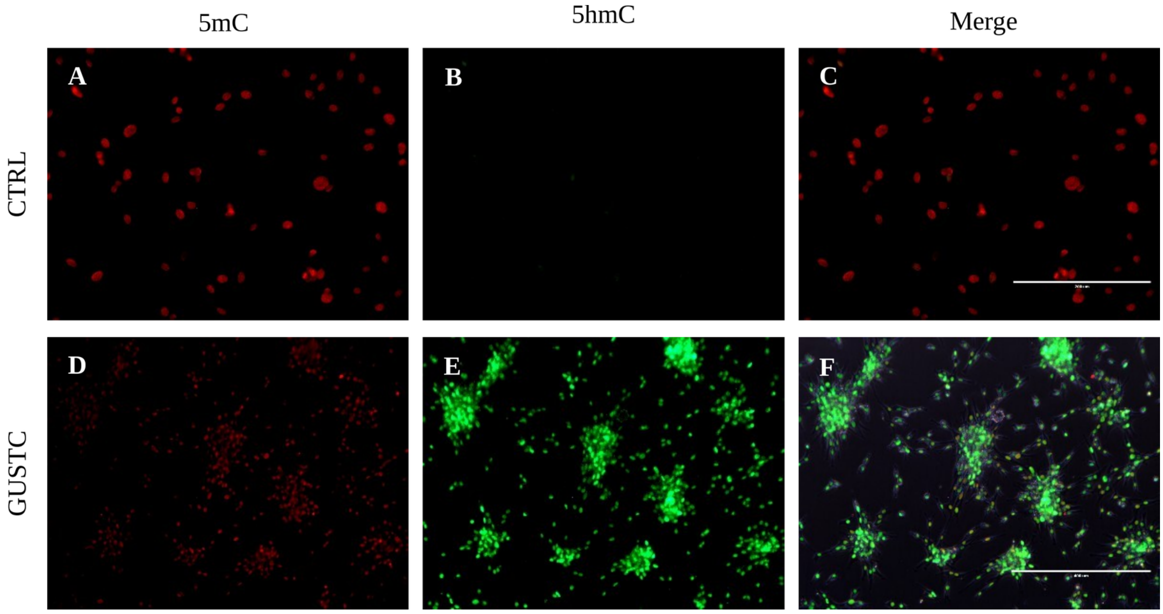

**Supplementary** Figure S12. Representative brightfield images illustrating the stepwise establishment of bovine embryonic stem cell (bESC) lines from nuclear transfer (NT)-derived blastocysts, from GUSTC- and control-treated donor cells. Day 8 blastocysts, blastocyst seeding onto MEF feeder layers, outgrowth at day 7, and an established bESC colony (white arrowhead) are shown for each group.

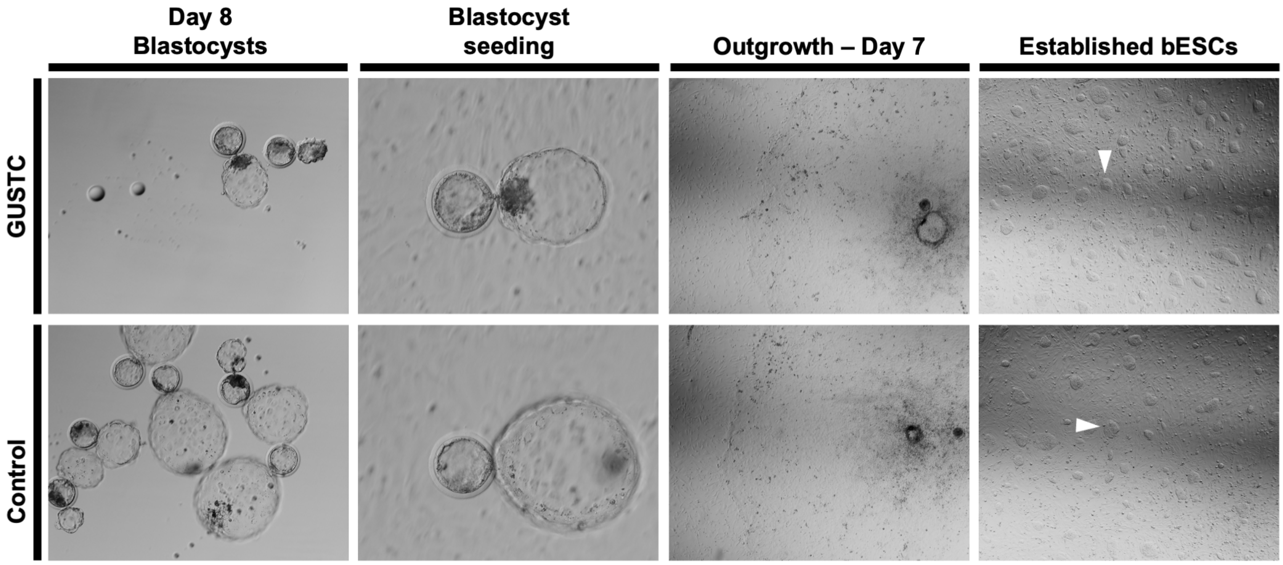
